# Strain-level diversity drives alternative community types in millimeter scale granular biofilms

**DOI:** 10.1101/280271

**Authors:** Gabriel E. Leventhal, Carles Boix, Urs Kuechler, Tim N. Enke, Elzbieta Sliwerska, Christof Holliger, Otto X. Cordero

**Affiliations:** Dept. of Civil and Environmental Engineering, Massachusetts Institute of Technology (MIT), Cambridge, MA, U.S.A.; Dept. of Environmental System Sciences, ETH Zurich, Zurich, Switzerland; Laboratory for Environmental Biotechnology, École Polytechnique Fédéral de Lausanne (EPFL), Lausanne, Switzerland

**Keywords:** Microbial communities, Strain diversity, Alternative states, Activated granular sludge, Accumulibacter, EBPR, Co-evolution

## Abstract

Microbial communities are often highly diverse in their composition, both at the level of coarse-grained taxa such as genera as well as at the level of strains within species. This variability can be driven by both extrinsic factors like temperature, pH, etc., as well as by intrinsic ones, such as demographic fluctuations or ecological interactions. The relative contributions of these factors and the taxonomic level at which they influence community structure remain poorly understood, in part because of the difficulty of identifying true community replicates assembled under the same environmental parameters. Here, we address this problem using an activated granular sludge reactor in which millimeter scale biofilm granules represent true community replicates whose differences in composition are expected to be driven primarily by biotic factors. Using 142 shotgun metagenomes of single biofilm granules we found that, at the commonly used genus-level resolution, community replicates varied much more in their composition than would be expected from neutral assembly processes. This variation, however, did not translate into any clear partitioning into discrete community types, i.e. the equivalent of enterotypes in the human gut. However, a strong partition into community types did emerge at the strain level for the most abundant organism: strains of Candidatus Accumulibacter that coexisted in the metacommunity--i.e. the reactor--excluded each other within community replicates. Single-granule communities maintained a significant lineage structure, whereby the strain phylogeny of Accumulibacter correlated with the overall species composition of the community, indicating high potential for co-diversification among species and communities. Our results suggest that due to the high functional redundancy and competition between close relatives, alternative community types are most likely observed at the level of recently differentiated genotypes but not higher orders of genetic resolution.

---

Microbes outside the lab rarely occur as individual clonal units, but exist as populations that are diverse on at least two levels: one, on the level of member species within microbial communities^1^; and two, on the level of genotypes within an individual ‘species’^2,3^. Recently, puzzling patterns have emerged related to both of these levels of diversity. Samples from similar environments have been found to differ greatly in their species-level composition, despite a high degree of conservation in community-level metabolic function^4,5^. Compositional differences manifest in terms of variation in species abundances that cannot be explained with simple models of neutral assembly from common species pools^6,7^. In this framework, communities are conceived as ‘patches’ that are colonized from a stable species pool. Explaining the large compositional differences generally requires assuming that dispersal is limited or that interspecific interactions determine which bacteria will dominate within a patch^8,9^.

In the extreme case, the processes of assembly, dispersal, and interactions can lead to the emergence of community types, i.e. distinct assortments of species on patches that do not appear to inter-mix with one another^10^, potentially due to incompatibilities between species. Concomitant with the variation in species abundance across patches, a more fine-grained assessment of genetic diversity reveals that what is commonly considered an individual species is often in fact a large collection of highly diversified genotypes. These genotypes combined can contain hundreds or even thousands of genes that are unique to only a subset of individual genotypes within the ‘species’^11–13^. Such nested taxonomic variability—with species abundances fluctuating strongly among replicate communities and each species itself acting as a ‘community’ of strain variants (i.e. a biological population)—is frequently overlooked in surveys of community structure due to the limitations inherent to the use of molecular markers (but see refs # [14–16] for some exceptions). This begs the question as to whether and to what degree strain-level variation influences community assembly processes, and whether the observed between-community compositional variation is not just a direct consequence of the within-species variation. Answering these questions requires (i) detailed knowledge of how much compositional variation remains when all environmental influences are accounted for, and (ii) how much strain-level diversity actually exists within each replicate community, where ecological interactions are frequent and can impact community assembly.

Detecting the effects of ecological interactions on community assembly is often difficult as it requires either strong external environmental consistency between sampled populations, or otherwise complete information of all environmental factors that are important for community function. Obtaining complete environmental information is generally impossible to date, as we lack the necessary understanding of the joint metabolic program of a whole community. Furthermore, any two natural communities will generally not experience uniform environments across the scales that are relevant for microbial interactions. Examples include communities around marine particles^17^, within soil aggregates^18^ or in dental plaque biofilms^19^. Full experimental control of the environment is in principle conceivable in a laboratory setting, but any experimental setup would need to be able to ‘grow’ communities to the level of complexity and diversity that adequately mimics natural communities. The lack of good environmental data at length scales that are relevant to the size of a community^20^ make the analysis of community structure and in particular community types extremely challenging, as apparent community types may just be the consequence of environmental variation unaccounted for during sampling.

In this paper, we use a well-controlled but complex engineered microbial community to ask: (i) how much residual community variation remains at relevant community scales when the external environment is uniform; and (ii) how strain-level variation structures microbial communities. In particular, we study the microbial community that constitutes the granular biomass in a laboratory scale granular enhanced phosphorus removal (EBPR) reactor^21,22^. EBPR is one of the prime forms of microbe-mediated wastewater treatment, as well as one of the best-studied model systems in microbial ecology. EBPR reactors have characteristic microbial communities depending on their operating conditions, often either enriching for phosphorus accumulating organisms (PAOs) or glycogen accumulating organisms (GAOs). Genomic studies of the microbial community in EBPR reactors have revealed that *Candidatus* Accumulibacter phosphatis (henceforth Accumulibacter) is the numerically dominant PAO^23,24^, and *Candidatus* Competibacter (henceforth Competibacter) is the dominant GAO, under respective favorable conditions^25–27.^ The Accumulibacter clade of uncultured organisms has been considerably well studied. Accumulibacter is actively involved in phosphorus removal by accumulation of intracellular polyphosphate, the primary sink of phosphorus in activated sludge^23^. Natural populations of Accumulibacter can be extremely diverse and reactor conditions select for a few specific micro-clades, leading to the establishment of a few dominant Accumulibacter variants within the reactor^28^,29. These micro-clades co-occur with other microbial taxa of diverse metabolic capabilities such as nitrifiers, denitrifiers or degraders of aromatic compounds^25^,30. Despite this diversity, EBPR reactors can maintain a relative stable performance for many months or even years.

The granular nature of the activated sludge biomass within bubble-column sequencing batch reactors imposes an interesting and unique spatial structure to the community: each granule can be seen as a selforganized replicate community within a well-mixed reactor environment. Thus, from an ecological standpoint, these reactors have two characteristic length scales: (i) the global reactor scale (meters), which represents a meta-community composed of multiple granule; and (ii) the local granule scale (millimeters), which corresponds to single replicate communities composed of many populations^31^. Functional consistency is generally assessed on the reactor scale—a length scale that is comparable to typical studies on marine or soil communities. However, biotic interactions such as cross-feeding, public goods exploitation^32^, antagonism^33^, and predation occur at the local single-granule scale and will likely control species abundances.

## Results

### Large between-granule variation within a single reactor

Our first aim was to determine the variation in microbial community composition across granules, or patches. To this end we sampled a total of 142 granules from a single experimental EBPR column reactor—the controlled metacommunity—at two time points (see Fig. 1) and performed shotgun metagenomic sequencing on individual granules (see Methods). We then mapped the sequencing reads to a database of reference genomes to obtain a genus-level community composition of each individual granule. Overall, 42,777,533 reads out of a total of 57,896,625 (74%) mapped to reference genomes in the database, of which 24,994,161 (43%) had normalized mapping quality of at least –1.86 (roughly 70% sequence identity) and were thus considered in the downstream analysis (see Methods and Figs. S1 and S2).

**Figure 1:**
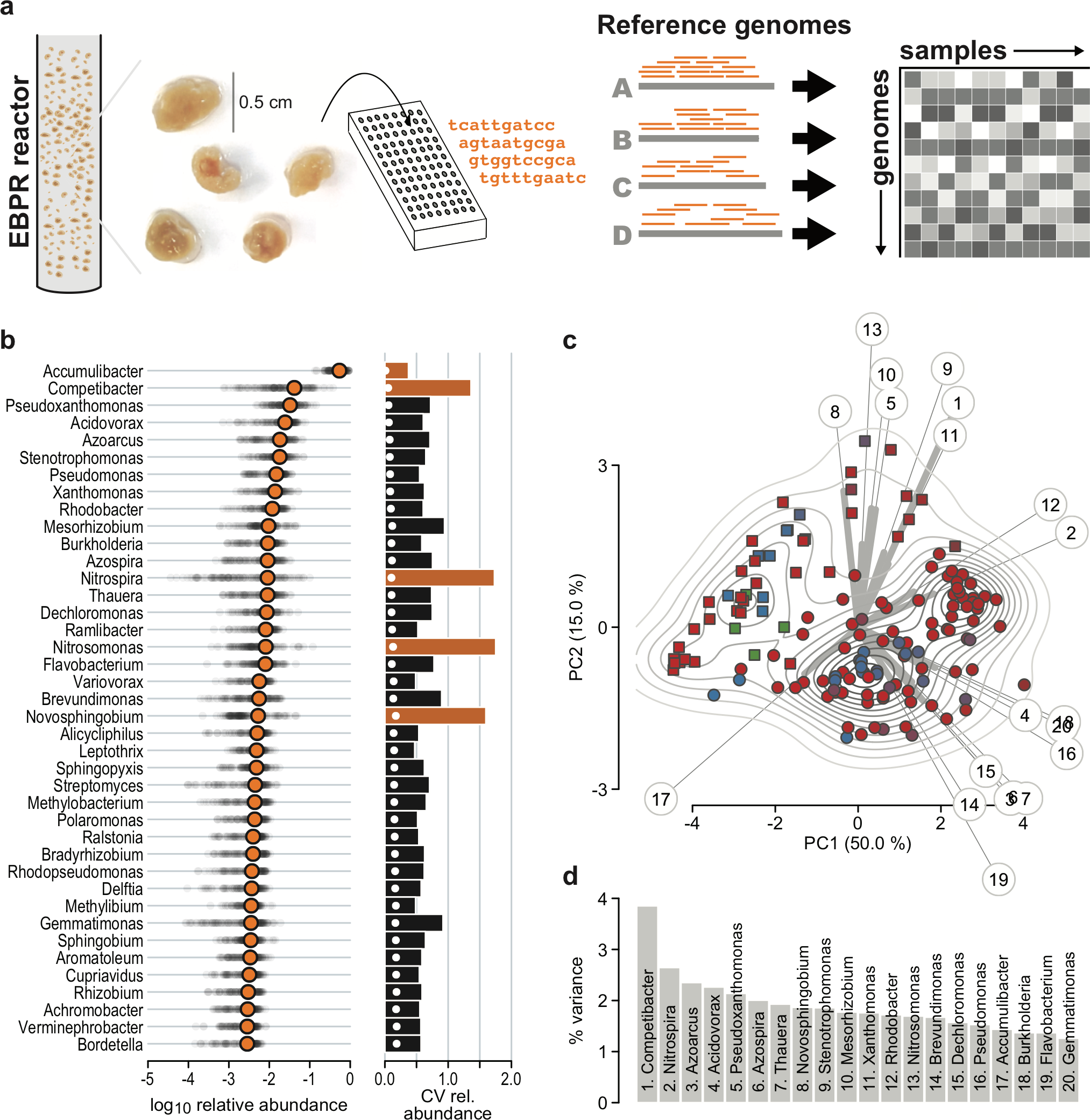
High degree of variability among replicate communities at the level of genera. **a.** Overview of the analysis process. Individual granules were sampled from an EPBR column reactor and submitted for shotgun metagenomic sequencing. Reads were then mapped to a database of reference genomes. The number of hits were normalized by genome length and tabulated by granule. **b.** Relative abundance of the top forty genera. Orange circles show the log^10^ mean relative abundance across all granules. The grey dots indicate the relative abundance of each genus per granule. The bars on the right show the coefficient of variation of the relative abundance for each granule.**c** Principal component analysis (PCA) projection of the log^10^ relative abundance variation across granules. The contour lines show a 2D kernel density estimate in the space of the first two principal components (PC). The grey lines indicate the direction of variation of the top 20 individual genera that explain the most variance, and the length of the thick portion indicates the amount of variation explained in the first two PC. The identity of genera is indicated by numbers corresponding to panel **d**. Granules were sampled in two batches (squares: May; circles: November), and the colors of the granules indicate the dominant Accumulibacter type (red: UW-1, blue: UW-2, green: BA-91). **d.** Amount of variance in the first two PC explained by the top 20 genera. The numbers correspond to the numbers in panel **c**.

The overall reactor-level composition was dominated by a single genus, *Candidatus* Accumulibacter, with a mean relative abundance of 54% (Fig. 1b). In total, 217 distinct genera recruited reads (Fig. S4), of which 47 comprised 90% of all reads. Within this set, the other dominant genera next to Accumulibacter were *Candidatus* Competibacter, *Acidovorax, Azoarcus*, and *Nitrospira*, which are typically found in these type of reactors^21^,26.

The reactor-level compositional consistency with previous studies, however, did not translate to uniform community composition in individual granules within a single reactor (Fig. 1b). Accumulibacter relative abundance fluctuated greatly across granules: from as low as 13% to as high as 93%. This large spread in relative abundance is captured by the coefficient of variation across samples, which was 37% for Accumulibacter. The coefficient of variation is the standard error divided by the mean and accounts for decreasing standard error as the relative abundance decreases. Despite this variation in relative abundance, Accumulibacter was rather conserved in comparison to the typical genus (Fig. S5). Competibacter, *Nitrospira, Nitrosomonas*, and *Novosphingobium* varied even more strongly than the typical genus, with coefficients of variation greater than 100% and overall variation of over two orders of magnitude (Figs. 1b and S5).

The variation in relative abundances was higher than expected under a model of neutral stochastic assembly from a single global species pool (i.e. metacommunity). A maximum likelihood fit of the relative abundances to a neutral model in the Sloan *et al.*^34^ framework predicts a very small value for the product of number of individuals per patch (granule) times the dispersal probability, Nm ˜358.4. Because the number of bacterial cells in a granule is much larger than this number, the dispersal probability under the neutral model would have to be very small to be consistent with the model predictions. Yet, even with such a small dispersal probability, the neutral model would predict a variation in relative abundance that does not match our data (see Fig. S6), and overall we reject the neutral model with *p <* 0.001.

Nevertheless, the large variability of specific genera across replicate communities within the same abiotic context raises the question of whether the granules might assemble into distinct community types, with neutral stochastic fluctuations around the typical composition of each type. Community types could then be interpreted as distinct species pools from which the replicate communities assemble.

A principal component analysis (PCA) of the log relative abundances only weakly partitions the granules into types (Fig. 1c). Granules mostly segregate according to sampling time (squares and circles in Fig. 1c), though the separation between time points is not perfect. No natural cutoff for the partitioning of within and between group variance appears in a *k*-means clustering (Fig. S7). Alternative ordination methods based on Bray-Curtis and Aitchison distances qualitatively reproduce this apparent lack of structure (Fig. S8).

The first two principal components jointly explain a large 55.7% of the overall compositional variation. We can extract and visualize the contributions from each genus to the variation explained by the principal components (grey rays in Fig. 1c). Interestingly, Accumulibacter variation is not the main explanatory factor for overall compositional variation (direction 17 in Fig. 1c). Granules are rather most strongly differentiated based on the abundance of Competibacter, followed by *Nitrospira*. Certain genera appear to covary in similar directions (e.g. 3, 6, 7; Fig. S9), and a portion of these appear to be conditionally dependent on each other (Fig. S10). Taken together, there is remarkable granule-to-granule variation in community composition within the reactor, yet no clear compositional classification into types emerges on the level of genera.

To test whether some of the weak structure found at the level of genera could be determined by microscale differences in abiotic parameters, we determined the effect of granule size on community structure, as size can have important consequences for the micro-scale environment, e.g. by limiting the diffusion of oxygen to the internal parts of the biofilm granule. We measured the size of the November 2014 granules, which had a mean diameter of 3.2 mm (*s* = 1.1 mm), where the smallest and largest granule had a diameter of 1.6 mm and 7.8 mm, respectively. Of the top eleven principal components that explain over 95% of the compositional variance, only principal components PC2, PC3 and PC11 correlated significantly with granules size after Benjamini-Hochberg corrections (see Tab. 1 and Fig. S11). This indicated that community composition did vary predictably with granule size in a compound manner along principal components PC2 (15.0%) and PC3 (10.0%), especially (PC11 only explained 0.75% of the overall variance). In order to visualize how individual genera change in relative abundance with respect to granule size, we calculated Spearman’s correlation coefficients for the top twenty most important genera in the PC2/PC3 direction. Among these that varied with granule size, potential denitrifiers were enriched in larger granules, while nitrifiers showed an enriched trend in smaller granules (Fig. S12). Hence, some of the weak structure observed at the level of genera can be partially explained by differences in granule size with respect to oxygen flow.

### Granules are differentiated based on strain-level differences

We hypothesized that the lack of clear differentiation between granules might also in part be due to the rather coarse grouping of genotypes based on genus. Although two populations of the same genus can have relatively similar function in terms of core metabolism (e.g. preferred electron donors and acceptors), even two genotypes within the same species can differ strongly in their ecological interactions, in particular those mediated by secondary metabolism^35^. Hence, the coarse-grained classification afforded by the genomic approach taken so far could obscure important patterns of ecological structure in the community. We tested this hypothesis by reconstructing the strain-level variation of Accumulibacter—the dominant species in the granules—which is known to display a remarkable genotypic diversity across reactors of different types and geographical location^36^.

Using a database of ten previously assembled partial and complete reference genomes of Accumulibacter, we found that at least three distinct genotypes coexisted within the reactor granular biomass (Fig. 2a). This was consistent with a classification based solely on the polyphosphate kinase *ppk1* gene, the most commonly studied marker of Accumulibacter diversity (Fig. S13a)^28,37,38^. Remarkably, these three genotypes excluded each other at the single granule level (Fig. 2a). Of the ten genomes assembled in previous surveys of Accumulibacter diversity, only the four genomes UW-1 (*ppk1* type IIA), UW-2 (IA), BA-93 (IA), and BA-91 (IIC) occurred at non-negligible abundances in our reactor, whereby UW-2 and BA-93 have been previously shown to be almost identical in their core genome^36^. For most granules—if UW-2 and BA-93 are considered a single reference—sequencing reads were almost exclusively recruited to one of the three reference genomes, respectively (Figs. 2a and S14). This implied the existence of alternative community types which are visible only at the level of strain variants. Given these observations we asked how much of the previously observed compositional variation across community replicates was driven by the three different Accumulibacter types that dominate the reactor. To test this, we first classified granules that consisted of 80% or more of a single genotype as ‘pure’ in terms of genotype. With this classification, 101 of the granules were of type UW-1, 22 of type UW-2 (or BA-93), 4 of type BA-91, and 15 were of mixed type UW-1 and UW-2. We then compared the compositional variation within and between ‘pure’ granules grouped by Accumulibacter type to ask to what extent Accumulibacter type conditioned the large variability previously observed. Overall community composition based on the top three principal components differed between granules of the same Accumulibacter type (ANOVA: *F* (2, 105) = 6.95*, p* = 9.41 · 10^−7^), indicating that some of the compositional variation at the community level is statistically linked to the type of Accumulibacter that dominated the granule. However, the significant differentiation was primarily driven by the different community composition observed in the four granules of type BA-91 (after removing four BA-91 granules, ANOVA: *F* (1, 102) = 1.94*, p* = 0.127; see also Fig. S8).

**Figure 2:**
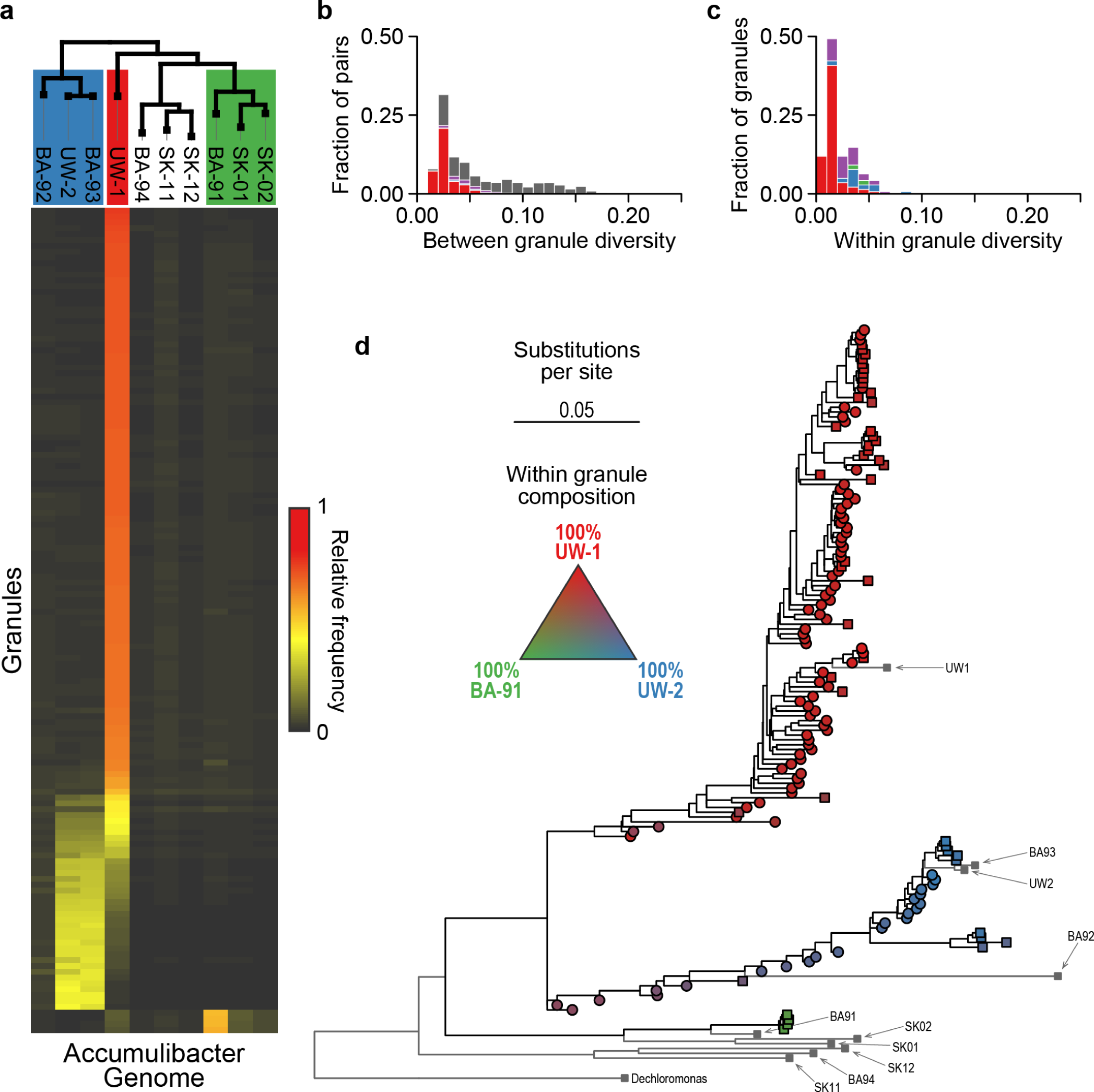
Strains of Accumulibacter segregate among community replicates. **a.** Heatmap of the relative proportions of reads that map to the 10 published reference genomes of Accumulibacter. Reference genomes are arranged in columns and individual granules are in rows, sorted by majority type for readability. The tree showing the relationship between reference genomes is taken from Skennerton et al.^36^. **b,c.** Histogram of the Accumulibacter nucleotide diversity between and within granules. The overall bar height indicates the total number/pairs of granules within the respective histogram bin, and the colors indicate the partitioning of that bin by majority Accumulibacter type. The color code follows panel **a**, with purple indicating granules of mixed UW-1/UW-2 type. **d.** Accumulibacter phylogeny of granules. Each tip corresponds to a Accumulibacter consensus sequence from an individual granule. The ten reference genomes from panel **a** are included in the tree and show the partitioning of granules based on type. The color gradient of the tips indicates the composition of granules from panel **a**.

An analysis of single nucleotide polymorphism (SNP) variation within and between granules confirmed that Accumulibacter was more clonal within granules than expected from the overall reactor diversity (Fig. 2b,c). To assess genetic diversity, we realigned all reads that were recruited to any of the Accumulibacter reference genomes by using a single reference genome as an alignment scaffold. For each aligned site we then calculated the nucleotide diversity, which is the probability that two randomly chosen individuals from within a granule will have a polymorphism at the given site, and averaged across the whole genome. Using this approach we estimated diversity both within single granules and between any two granules. Fig 2b shows that the distribution of between-granule diversities is characterized by a long tail corresponding to the inter-type comparisons (e.g. type UW-1 *vs.* UW-2), which is absent from the single granule diversity distribution (Fig 2c). Moreover, diversity between granules of the same type (e.g. red bars in Fig 2c for UW-1) was still higher between granules than within them, indicating that each Accumulibacter type carried significant strain-level diversity that segregated across, more than within granules. This observations made us question whether the micro-evolutionary history of Accumulibacter beyond the classification into three broad types could be linked to the variability observed at the community level.

### Lineage-level genetic structure correlates with compositional structure and shows signs of codiversification

In order to study the micro-evolutionary history of Accumulibacter within the granular biomass of the reactor, we took advantage of the comparatively low genomic Accumulibacter diversity within granules. We reconstructed three phylogenetic trees from consensus sequences based on realignment against each of the three reference genomes, and combined them into a single phylogeny of Accumulibacter across granules (see Fig. S15), where each tip of the tree represents the Accumulibacter population in a given granule. In agreement with our previous analysis, the phylogenetic structure reflected the large distances between Accu mulibacter types, with the UW-1, UW-2, and BA-91 granules segregating in different clades (Fig. 2d).

The phylogeny also revealed a more fine-grained association between the Accumulibacter genetic structure and the ‘auxiliary’ community composition (i.e. excluding Accumulibacter). The matrix of phylogenetic distances between granules correlated significantly with compositional distance (number of tips, *n* = 142; Spearman’s *ρ* = 0.149; Mantel test *p* < 0.001, 1000 permutations). There was no significant correlation between the Accumulibacter genetic structure and the relative abundance of Accumulibacter only (number of tips, *n* = 142; Spearman’s *ρ* = 0.0497; Mantel test *p* = 0.071, 1000 permutations), indicating that Accumulibacter genetic structure correlates with its auxiliary community rather than Accumulibacter abundance itself. This association was only in part driven by the difference in composition of granules of type BA-91, as the correlation persisted even when those granules were excluded from the analysis (*n* = 138; Spearman’s *ρ* = 0.122; Mantel test *p* < 0.001, 1000 permutations). The correlation can also be inspected visually, where sub-clades of the Accumulibacter tree—i.e. granules with genetically similar Accumulibacter populations— are also more similar in the overall community composition (Fig. 3). Hence, the fine-scale genetic lineage structure of the Accumulibacter populations in each granule corresponds to the fine-scale differences in the composition of the auxiliary community to Accumulibacter.

**Figure 3:**
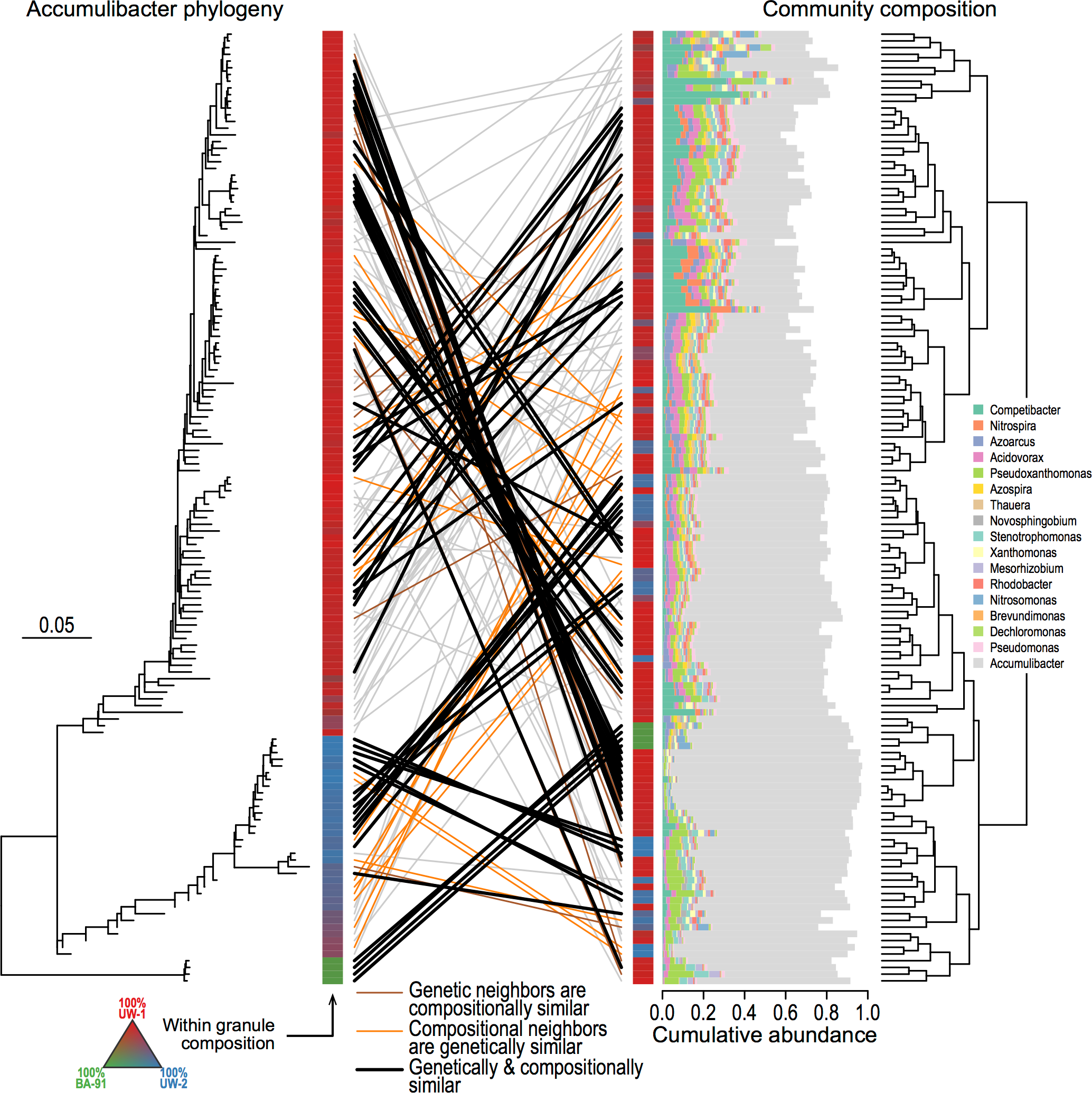
Accumulibacter strain-level phylogeny correlates with compositional structure. Left: The phylogeny is the Accumulibacter phylogeny from Fig. 2c. Right: The composition of each granule is shown using the relative abundances of the top 10 genera from Fig. 1b. Granules are ordered according to a hierarchical clustering based on the Aitchison distance. The tips of the phylogeny (left) are connected to the corresponding compositional bar (right) by lines. Lines are colored according to the following criterion: orange, the genetically closest neighbors of the granule in the phylogeny are more compositionally similar that would be expected by chance; brown, the compositionally closest neighbors are genetically more similar than would be expected by chance; black, both is true; grey, no significant association.

Although ecological associations between taxa are often studied at the genus or species level, there is *a priori* no reason why interactions should be confined to those levels of genetic resolution. Informed by the link between the population genetic structure of Accumulibacter and the community composition, we extended the phylogenetic analysis to other genera to ask whether associations between different taxa occur at the strain-level. Similarly to Accumulibacter, we determined SNP frequencies with a coverage of at least three and reconstructed phylogenies from consensus alignments for eight of the most variable genera across samples: Competibacter, *Dechloromonas, Acidovorax, Nitrospira, Nitrosomonas, Thauera*, and *Azoarcus*. The genetic structure of the individual genera correlated significantly across granules (Fig. S16). This pattern suggests that some bacterial populations are co-diversifying within the reactor environment.

## Discussion

This paper uses the granular biofilm communities in an activated granular sludge reactor as a controlled model system with which to tackle a fundamental problem in microbial community ecology and evolution, namely the interplay between community structure and strain-level variation in bacteria. First, we found that community composition at the level of genera is highly variable across replicate communities—i.e. granules— despite a homogeneous external abiotic environment and a common inoculum (i.e. global species pool). Second, while individual replicate communities are only poorly distinguished on the basis of genus-level taxonomic characterization, strain-level genetic resolution of member species reveals a clear partitioning of replicates into community types. Third, communities that are genetically similar in the abundance of the dominant member Accumulibacter are also more similar in overall community composition, and there is indication that certain member populations display similar evolutionary histories across granules.

The surprising large variability in community composition indicates to us that biological interactions play a significant role in the assembly of the local patch communities in granules. The amount of variation that would be expected from neutrally assembling microbial communities has often been viewed in the light of ecological theory of macro-organisms, such as mainland-island biogeography and Hubbell’s Unified Neutral Theory of Biodiversity. These theories predict that local island or patch communities can indeed vary in their composition if either the mainland/meta communities from which they are colonized are different, or if dispersal between patches is small enough such that initial differences in colonization cannot be equalized. For microbes, however, the question of dispersal ability has mostly been discussed at length scales that are orders of magnitude larger than an individual cell^39^. In our model system, there is strong periodic mixing such that the bacteria should in principle not be limited by dispersal between patches^21^. Therefore, we posit that the large variability in relative abundances between communities is due to interspecies interactions that can preclude or facilitate the coexistence of specific community members. For example, the *Candidatus* Competibacter clade varies over multiple orders of magnitude across granules, and—importantly—is strongly anti-correlated with Accumulibacter. Previous studies have shown that Accumulibacter and Competibacter are in direct competition and can be preferentially enriched for depending on operating conditions^27^. The reactor studied here was fed with both propionate and acetate as a carbon source. Propionate-only reactor conditions can provide Accumulibacter with an advantage over Competibacter, while acetate-only reactors have previously resulted in more erratic compositional dynamics with Competibacter dominating at times^25,27^. Given this differential effect of carbon source on the dominant species at the reactor scale, it is intriguing to consider the coexistance of Accumulibacter and Competibacter in terms of niche partitioning at the granule scale. Such niche partitioning is generally employed to explain the coexistence of two or more competitors that occupy the same habitat^40,41^. In the case of our granules, while the availability of two carbon sources might indeed stabilize the coexistence of Accumulibacter and Competibacter on the metacommunity scale, they nevertheless do appear to segregate on the individual patch scale, despite the *a priori* availability of niches on each granule. While the respective preferred carbon source of either organism should be homogeneously distributed across the whole reactor, it is likely not the only resource for which they compete. This highlights that—at least for bacteria—niche differentiation can rarely be reduced to a single nutrient dimension.

The differentiation of replicate communities on the basis of Accumulibacter type highlights the importance of phylogenetic resolution when studying interactions and their effect patterns of community structure. At coarse-grained levels of genetic resolution—such as the level of genera—the patterns of community structure are weak and communities cannot be clearly partitioned into discrete types (Fig. 4). Interestingly, most other comparative studies of microbial communities, for example of the human gut microbiome, are also based on coarse-grained phylogenetic markers like 16S rRNA and similarly detect only weak patterns of structure^42^. In contrast, our study is to our knowledge one of the first that attempts to compare communities at deep genetic resolution and does indeed find a clear pattern of community partitioning due to the mutual exclusiveness of the various genotypes of the same species. Although such an extreme physical segregation between strains seems to indicate that each granule grows from a clonal colony of Accumulibacter, given the high density of bacteria in the reactor it is not obvious how such strong founder effects could be maintained in the absence of more direct competitive interactions. We therefore hypothesize that the observed patterns of exclusion between community types reflect strong competition between functionally equivalent classes in an environment with strong spatial structure.

**Figure 4:**
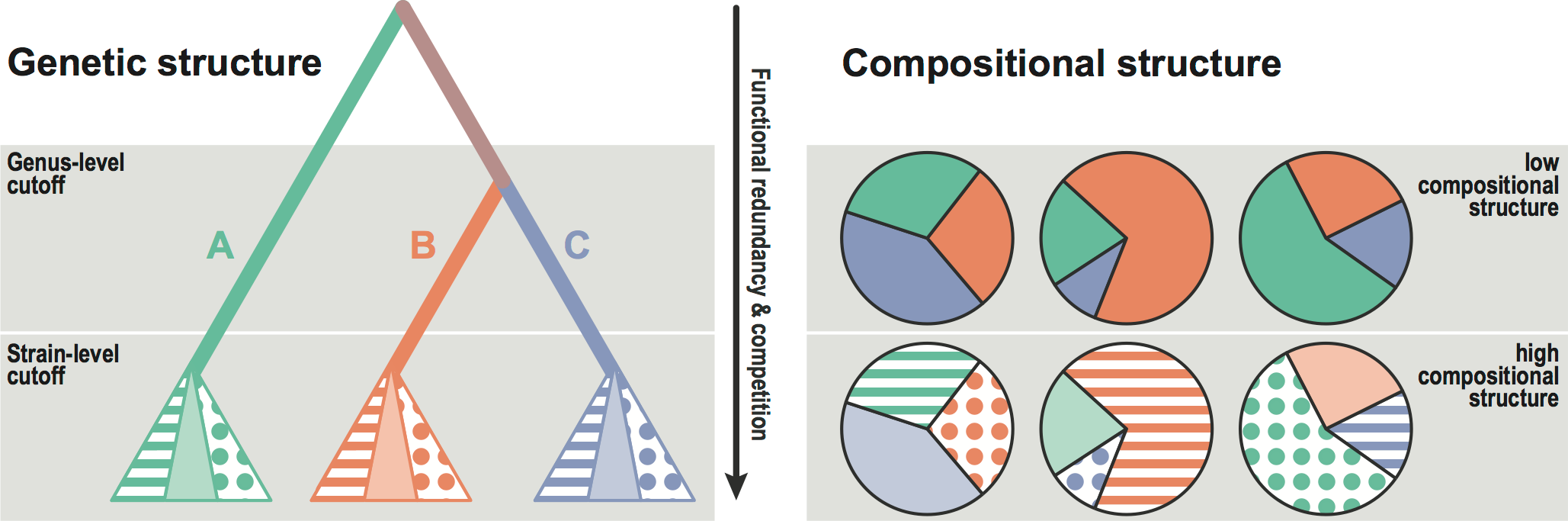
Schematic view of the relationship between phylogenetic depth and community structure. At the level of clades (e.g. genera) there is high variability among replicate communities but little structure in community space (no community types). However, closer to the tips of the phylogeny, at the level of strains within species, strong community types emerge. We propose this pattern emerges because of the high functional redundancy and competition among strains. These two features imply that two replicate communities can be functionally equivalent (to first approximation) independently of the strains that populate them, and that the co-occurrence of two ‘strains’ in the same granule is ecologically unstable and therefore unlikely to be found.

A crucial question is to what extent the three types of Accumulibacter are functionally distinct. Previous studies have investigated differences in Accumulibacter populations from different *ppk1* clades. One suggested functional differences is the potential ability of type IA to use nitrate as a terminal electron acceptor, while type IIA might not^36^,43,44. Unfortunately, using our genomic approach it is not straightforward to detect these particular differences, as the genomes of both types have been shown to contain the necessary genes, but it is still unclear whether the pathways are functional in both types. These and potentially other functional differences between Accumulibacter types could indeed result in differential attraction of auxiliary community members. This is in part supported by the correlation between the Accumulibacter genetic structure and the auxiliary community compositional structure. A deeper investigation into the distribution of functional genes across granules and the organisms that contain them in their genomes might prove an intriguing path forward.

The link between Accumulibacter genetic structure and community composition, as well as the apparent correlation between phylogenies of different genera provides a means for speculation about the life history of granules in a reactor. Two scenarios for the appearance of such signals are: (i) as a granule initially assembles and ‘grows’, the specific genetic variant of Accumulibacter in the granule determines both the relative abundance of the other members as well as specifically those genetic variants with which this particular Accumulibacter clone was previously associated in the past; (ii) the life history of granular biofilms involves cycles of growth, break-up or budding-off, and re-growth. Both scenarios would result in similar phylogenetic patterns, yet (i) requires preexisting co-adaptations between specific lineages while (ii) might just reflect correlated neutral diversification. As so often is the case, reality is likely a combination of both scenarios. Because the diversity found in the Accumulibacter phylogeny is most likely rather old and pre-dates the diversity generated in the reactor, we hypothesize that alternative community types—each containing a different Accumulibacter type—were formed early during the establishment of the granular sludge. These community types could then have been maintained by granule fission, and their frequencies regulated by competition between full communities, leading to the mixed granule assemblages found in extant reactors.

Because of the tight spatial linkage between organisms over ecological and evolutionary timescales, granular biomass offers a unique opportunity to study the drivers of microbial community structure and evolution. Full-granule fission—although not directly observed but only inferred in this paper—plays a crucial role in conferring cohesive evolutionary histories to granules, akin to evolutionary histories of individual organisms. Even though scenario (ii) above does not a priori call for lineage-level co-adaptations between organisms as a prerequisite, the spatial coherence does set the stage for co-evolution to occur.

Overall, the study of granular biomass communities that co-occur in a unified reactor environment illustrates a previously untapped resource to inform long-standing questions in microbial ecology. Granular communities allow for the necessary experimental control of the abiotic external environment, while conserving community differentiation at length scales that are relevant for microbial interactions^20^. Importantly, they provide an ideal platform to combine high-resolution genetic studies of individual species with community function and stability.

## Methods

### Experimental methods

#### Aerobic granular sludge reactor and sample collection

The aerobic granular sludge was cultivated in a double-wall glass bubble-column reactor (BCR) with a working volume of 1.75 L, an internal diameter of 62 mm, and a height-to-diameter ratio of 11. The synthetic influent wastewater contained equal concentrations of acetate and propionate as carbon and energy sources in terms of chemical oxygen demand (COD: 300 mg/L; 2.7 mM or 200 mg/L propionate, 4.7 mM or 280 mg/L acetate). Ammonium and phosphate were the main nutrient sources. The remaining composition of the influent was the same as described in Lochmatter & Holliger^22^. The reactor was operated with a sequencing batch regime (SBR) and the fixed 3 h SBR cycles consisted of the following phases: anaerobic feeding without mixing (60 min), constant aeration (110 min), settling (5 min), and withdrawal (5 min; 50% of total volume). The hydraulic retention was thus 6 h. The biomass concentration and the sludge age were maintained at 15 ± 2 g_VSS_/L and 15–20 days, respectively, by automated purge of excess sludge from the aerated mixed liquors. Temperature was controlled at 20°C, and pH was regulated at 7.0 ± 0.2 by dosing HCl or NaOH at 0.5 mol/L. Samples from this reactor were collected once in May 2014 and a second time in November 2014 for manual sorting of individual granules and DNA extraction.

#### DNA extraction and sequencing

Genomic DNA was extracted from granules with the MasterPure DNA Purification Kit (Epicentre #MCD85201) following the standard protocol with an additional overnight enzymatic digestion step. Genomic libraries were prepared using the Nextera XT Library Prep Kit following the manufacturer’s protocol. The resulting mean fragment size was around 500bp. Sequencing was performed on Illumina MiSeq (250bp paired-end) at the Genomic Diversity Center (ETH Zurich, Zurich, Switzerland) and Illumina HiSeq (100bp paired-end) at the Whitehead Institute of Biomedical Research (MIT, Cambridge, MA, U.S.A.) for the May and November granules, respectively. The sequencing runs resulted in a total of 8,799,127 (46 samples) and 41,060,893 (96 samples) paired-end reads, respectively, for a total of 99.7M reads.

#### Determination of granule size

We determined granule size only for those granules from November 2014. Sizes of granules were measured by drawing a two dimensional polygon around the contour of imaged granules and calculating the enclosed area, *<u>A</u>*. The granule radius was approximated by assuming a perfect circle with area equal to the polygon, 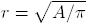.

### Statistical methods

#### Reference database construction

Raw sequencing reads were preliminarily run through MG-RAST^45^ to obtain a rough representation of genera contained in the reactor. Representative genomes of the genera that accounted for 90% of the mapped reads were then downloaded from NCBI and complemented with the Accumulibacter references genomes^36^ and the Competibacter genome^46^. A full list of accession numbers is given as Supplementary Data.

#### Quality trimming and merging

The paired raw sequencing reads were first merged and trimmed with PEAR 0.9.0^47^. Reads that were not merged were additionally quality trimmed with Trimmomatic 0.36^48^ (`LEADING:3, TRAILING:3 SLIDINGWINDOW:10:20, MINLEN:36`). Overall, this produced a data set of 57,896,625 reads.

#### Read mapping

All reads were mapped against the representative database using Bowtie2 2.3.3.1^49^. Because we expected the reference genomes in the database to be quite different from the actual genotypes in the reactor, we used end-to-end alignment with very relaxed settings and no minimum cutoff (`--score-min L,0,-10 -N 1`). This allowed us to obtain the best hit—if any—for each read and then subsequently filter out reads in a second step based on the alignment score using `bamcnt`^50^. We normalized the alignment scores from Bowtie2 by the read length, and fit a Gaussian mixture model to scores using the `R` package `Rmixmod`^51^. This revealed three classes of alignment scores (Fig. S1), which we roughly attributed to: (A) matches to a correct genotype, normalized alignment score > –0.36, which translates to 6 mismatches per 100 base pairs when there are no insertions/deletions; (B) matches to a related genome, normalized alignment score > –1.86, which translates to 31 mismatches per 100 base pairs without mapping gaps; and (C) matches to distantly related genomes or spurious matches. We subsequently only considered reads from categories A and B.

#### Genus 16S tree

For each genus in our reference database, we extracted the 16S sequence from the NCBI 16S Microbial Sequences database (Bioproject 33175; Aug 6, 2016). Sequences were aligned using Clustal Omega 1.2.14^52^, and the phylogeny was subsequently reconstructed with FastTree 2.1.9^53^ using a GTR model.

#### Genus abundances

The genus of each reference genome was obtained from the taxonomic classification in NBCI. Reads were then grouped and enumerated for each granule according to the genus of the matched reference genome. In order to avoid double counting of DNA fragments for paired reads, we only counted the match of the forward read. Reads that mapped to reference sequences that were identified as plasmids were discarded. Because the expected number of DNA fragments in the library is proportional to the overall genome size, we further normalized the genus read count by the average genome length (excluding plasmids) of each genus in the database. Relative abundances were calculated by dividing the normalized counts by their sum for each granule.

#### Coefficient of variation

We first calculated the coefficient of variation (CV) of the relative abundance, *r*_*ig*_, for each genus *g*: *v*_*g*_ = sd (*r*_*ig*_) /mean (*r*_*ig*_). Even if the composition of each granule were equal, we would expect a certain CV just because of random sampling from the population. Assuming that sampling is uniform with respect to each member, the distribution for the abundances should follow a multinomial distribution. Hence, the expected CV of the relative abundance just as a result of sampling is 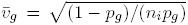, where *p*_*g*_ is the true relative abundance and *n*_*i*_ is the sampled population size. Using the number of reads as a proxy for the sampled population size, and choosing the lower bound of the number of reads across all granules as *n* = 10^4^ (see also Fig. S2), we obtain a conservative estimate for the expected CV purely due to sampling effects.

Additional noise from technical sources (DNA extraction, library preparation, etc.) might furthermore contribute to the observed variation in relative abundance. We thus derived a heuristic approach to identify genera that showed an especially large (or small) coefficient of variation. We first regressed the CV onto the log_10_ mean relative abundance of the genera. Because the amount of expected variation increases with decreasing relative abundance, we excluded a small number of genera with a particularly low mean relative abundance of below 10^−4^ from the regression (see Fig. S5). We then calculated Cook’s distance of the CV for each genus—a standard measure used to identify outliers in regression analysis. Genera that had a Cook’s distance of more than two standard deviations above (or below) the mean in log space were highlighted as strongly varying (or conserved).

#### Neutral model

We tested whether the observed relative genus abundances arose from a neutral community assembly process using the framework of Sloan *et al.*^34^. The Sloan model can be viewed as a large population size diffusion approximation of Hubbel’s Unified Neutral Theory of Biodiversity^*6*^. In absence of growth rate differences between the individual genera, the probability density that a genus, *g*, within an individual patch, *i*, has relative abundance *r*_*ig*_ depends on the relative abundance in the metacommunity, *p*_*g*_, the migration probability into the patch, *m*, and the total patch population size, *N*_*i*_, and follows a beta distribution,

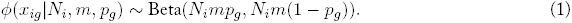

Note, that the parameters *N*_*i*_ and *m* only occur together as a compound parameter *m*_*i*_ = *N*_*i*_*m*. The overall log likelihood that our data arose from a neutral model then just is

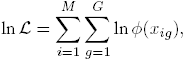

where *M* is the total number of patches (granules) and *G* is the number of genera. Assuming that the population sizes are the same across all patches, *m*_*i*_ = *m ∀i*, we find the maximum likelihood estimate of *m* and (*p*_1_*,…, p*_*G*_) by maximizing ln *ℒ*. Because of the constraint ∑_*i*_ *p*_*i*_ = 1, we use the transformation *y*_*i*_ = *p*_*i*_/*p*_1_ for *i* = 2*,…, G* and maximize the likelihood numerically using the subplex algorithm implemented in R^54^. The metacommunity relative abundances are easily found by back transforming the *y*_*i*_ using *p*_1_ =1/(1 +∑_*i*_ *y*_*i*_) and *p*_*i*_ = *y*_*i*_*p*_1_.

We assess the goodness of fit analogous to Etienne^55^, by simulating data sets under the maximum likelihood Sloan model and comparing the likelihoods of the simulated data to the real data. The number of simulated data sets that have a likelihood that is smaller than the real data yield empirical *p*-values for the support of the neutral model. Furthermore, the Sloan model yields an expectation for the mean and variance of relative abundances in each patch, and hence also for the coefficient of variation,

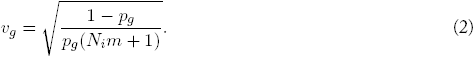

Note, that when migration/dispersal is frequent, *m →* 1 and the coefficient of variation tends to that of a multinomial sampling model 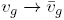.

#### Multivariate analysis

We performed a principal component analysis (PCA) by singular value decomposition of the sample covariance matrix of center log-ratio transformed (CLR) relative genus abundances^56^. A pseudo count equal to the mean genus count per sample was added to counts from each sample to account for uneven sequencing depths. Genus counts in each sample were then normalized by the total number of genus counts per sample. Genus rays were extracted as the eigenvector components of the top two principal components. Additionally, we also performed multidimensional scaling on Aitchison distance—Euclidian distance of CLR vectors—and Bray-Curtis dissimilarity matrices using `metaMDS` in the vegan package^57^. We assessed the partitioning of granule communities into discrete units through a *k*-means clustering of the CLR transformed values with cluster numbers of 1, 2,*…*, 30. Clustering at different cluster numbers was compared on the basis of the within-cluster sum of squares *versus* the overall sum of squares. We determined conditional independence between genera from the inverse covariance matrix of CLR transformed relative abundances. We estimated the inverse covariance matrix using a graphical LASSO regularization approach^58^ with a regularization parameter of *ρ* = 0.001.

#### Accumulibacter genomic diversity

In order to assess Accumulibacter genomic diversity, we first re-aligned all reads that mapped to any Accumulibacter reference genome to the UW-1 reference. For each base pair position on the genome that were covered by at least three reads, we then estimated allele frequencies, {*p*_*A*_, *p*_*C*_, *p*_*G*_, *p*_*T*_}, as the fraction of nucleotide variants that were observed. We calculated within granule nucleotide diversity as the probability that two randomly chosen DNA fragments would have different nucleotide variants at a base position *k*,

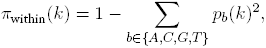

and averaged this over all sites with coverage of at least three using `vcf2hs`^50^. Similarly, we calculated nucleotide diversity between granules *i* and *j* using `vcf2fst`^50^ as the average across all sites *k* with coverage of at least three in both samples *i* and *j*:

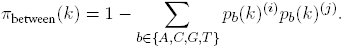

We then generated Accumulibacter consensus sequences for each granule using the majority nucleotide at each base with coverage of at least three using `vcfConsensus`^50^.

In order to reconstruct Accumulibacter phylogenies including the ten reference genomes, we additionally randomly fragmented all ten reference genomes as well as the *Dechloromonas* reference into 10,000 fragments of 1000 base pairs each using `fastaFrag`^50^. We aligned these to the UW-1 reference and called consensus sequences in the same manner as for the granule samples. Finally, we reconstructed phylogenies using FastTree 2.1.10^53^ with the gamma-optimized CAT model and the default pseudo count, and subsequently rooted the trees with Dechloromonas as an outgroup. To account for potential biases of using the UW-1 reference, we repeated this procedure for the UW-2 and BA-91 references. We calculated a combined cophenetic distance as the element-wise geometric mean of all three cophenetic distance matrices, and built a tree from the combined distance matrix using least-squares FastME^59^ implemented in ape^60^.

#### Comparing the Accumulibacter phylogeny and the community composition

We calculated the Spearman correlation between the combined cophenetic distance matrix and the community distance measures: (i) the Aitchison distance of the whole community; (ii) the renomalized subcommunity with Accumulibacter removed, and (iii) the distance between granules based on Accumulibacter relative abundance only. To test the overall significance of these correlations, we performed Mantel tests, which involves shuffling the rows and columns of the distance matrices, recalculating the correlation coefficient, and enumerating the number of times the shuffled correlation coefficient is larger than the correlation coefficient calculated from the full data.

To identify coherent subclades we calculated the geometric mean phylogenetic or Aitchison distance, respectively, of the three closest neighbors, and compared this to the geometric mean distance of 1000 randomly chosen triplets. A granule was marked as coherent if fewer than 5% of the random triplets had a geometric mean distance that were below the actual nearest neighbor geometric mean distance.

#### Comparing phylogenies between genera

For a subset of genera, we generated consensus alignments in the same manner as for Accumulibacter—i.e. by realigning all reads that mapped to the genus onto a single reference genome—and subsequently reconstructed phylogenies using FastTree (see Figs. S17, S18, S19). We used the following reference genomes: *Acidovoarx sp.* (NC_018708.1), *Azoarcus sp.* (NC_008702.1), *Candidatus* Competibacter denitificans (GCF_001051235.1), *Dechloromonas aromatica* (NC_007298.1), *Nitrosomonas sp.* (NC_015731.1), *Nitrospira defluvii* (NC_014355.1), *Thauera sp.* (NC_011662.2). Granules for which fewer than 1000 bases had a coverage of at least 3 were dropped from the trees. We calculated the Spearman correlation between the cophenetic distances for pairs of genera and assessed significance with Mantel tests.

#### Marker gene analysis

We performed a multiple sequence alignment of 1 012 published Accumulibacter *ppk1* sequences and a Rhodocyclus *ppk1* sequence as an outgroup using MUSCLE^61^. A phylogenetic backbone tree was then reconstructed using RAxML with GTRGAMMA rate variation. We identified 2 897 *ppk1* sequence reads using HMMER 3.1b2^62^ based on the 1 012 sequences reference set. These reads were then added to the multiple alignment with MUSCLE. Finally, the identified reads were phylogenetically placed on the backbone using pplacer 1.1.alpha17^63^.

## Acknowledgements

We thank Tomas de Wouters and Seppe Kuehn for comments on an early draft of the paper. GEL was supported by the Swiss National Science Froundation (162251) and the Human Frontiers Science Program (LT000643/2016-L). OXC was supported by a grant from the Simons Foundation (542395).

## Supplementary Information

**Table 1:** Principal compontents PC2 and PC3 that explain an important proportion of the variance correlate significantly with granule size.

| | % variance | $S$ | $\rho$ | $p$ (adjusted) |
| --- | --- | --- | --- | --- |
| PC1 | 50.0% | 138342 | 0.062 | 1 |
| PC2 | 15.1% | 82281 | 0.442 | $7.21 \cdot 10^{-5}$ |
| PC3 | 10.0% | 210504 | -0.428 | $1.52 \cdot 10^{-4}$ |
| PC4 | 8.44% | 176522 | -0.197 | 0.595 |
| PC5 | 3.13% | 157249 | -0.067 | 1 |
| PC6 | 2.39% | 136372 | 0.075 | 1 |
| PC7 | 2.27% | 153590 | -0.042 | 1 |
| PC8 | 1.38% | 146497 | 0.006 | 1 |
| PC9 | 1.00% | 142847 | 0.031 | 1 |
| PC10 | 0.97% | 138705 | 0.059 | 1 |
| PC11 | 0.75% | 92748 | 0.371 | $2.19 \cdot 10^{-3}$ |

Spearman's rank correlation of granule size was calculated for the top 11 principal components that account for 95% of the variance. The reported $p$ -values for a correlation different from zero are Hochberg corrected.

**Figure S1:**
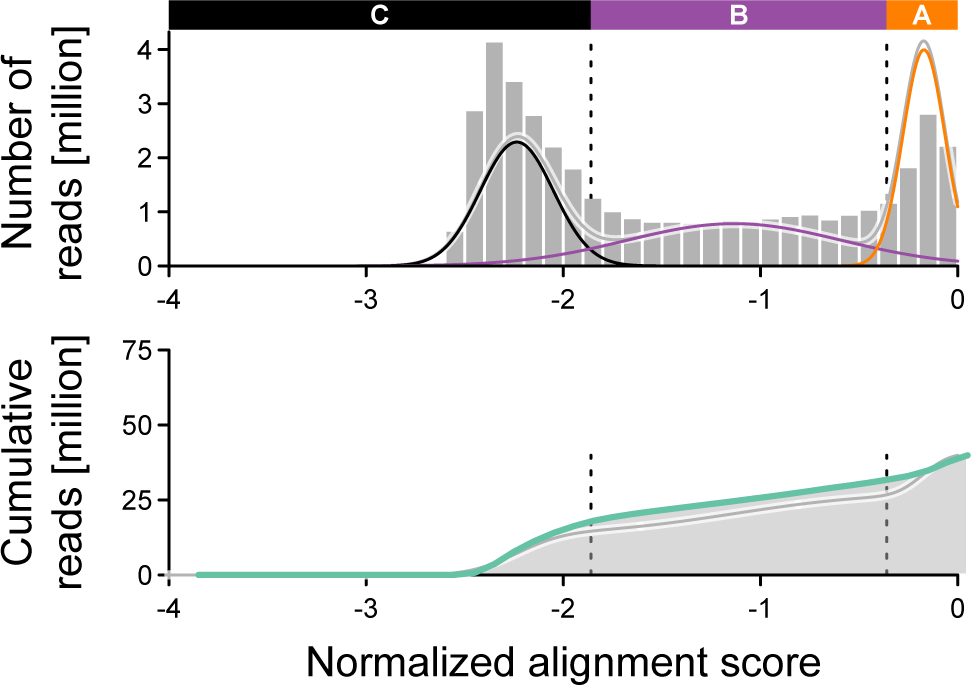
Distribution of alignment scores for reads. Alignment scores are calculated using the default scoring from Bowtie2^49^: mismatches are penalized with a score of 6, gap openings by 5 and gap extensions by 3. Scores are then normalized by read length. A read score of 1.0 thus implies a mismatch at every sixth base on average (if there are no gaps). We identified three regimes of score matching: (A) strong match, likely a correct reference genome for the specific genome in the sample; (B) weak match, likely a close reference genome of the species or genus; and (C) spurious matches. All analysis in the main text was performed using a cutoff of 1.86 that includes regimes A and B.

**Figure S2:**
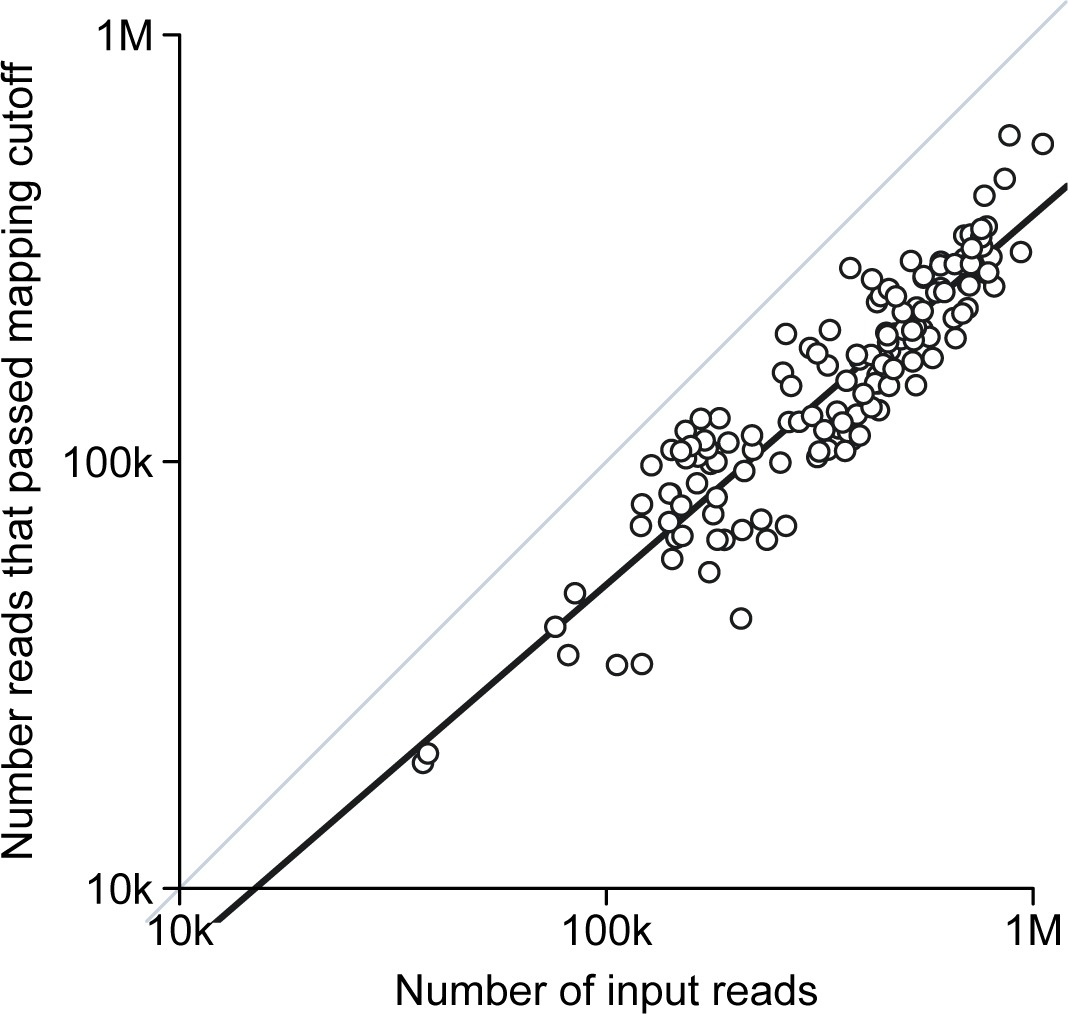
Number of reads that passed the mapping cutoff as a function of the total number of sequencing reads per granule. The number of reads that mapped to an entry in the reference database with sufficient normalized score (*>* − 1.86) did not differ systematically between granules.

**Figure S3:**
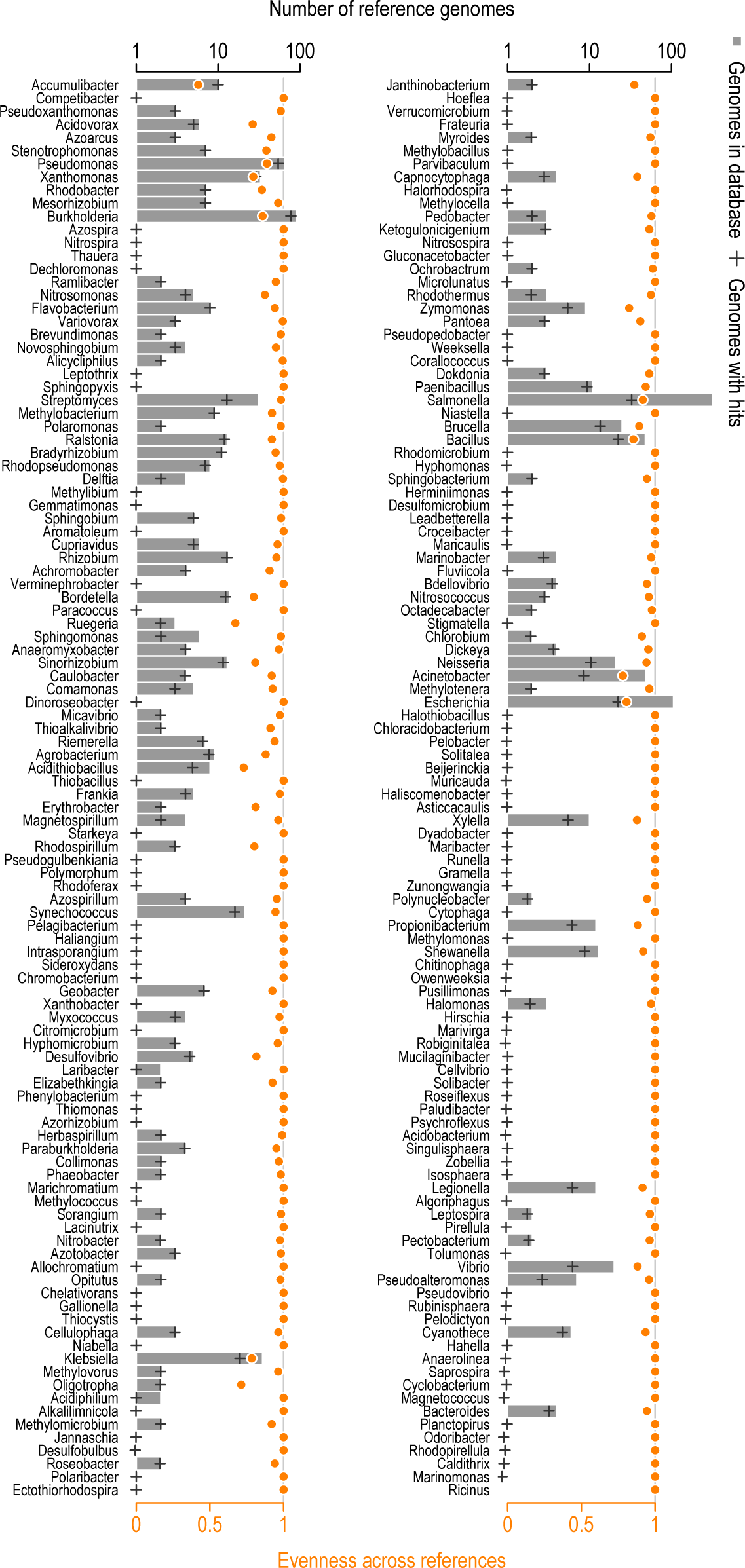
Reference genomes per genus. The grey bars show the total number of reference genomes per genus in our database. Genomes with multiple chromosomes/contigs are grouped by NCBI taxonomic ID. Black + show the average number of taxa across samples that recruited reads. Orange points show the evenness of recruitment across taxa. An evenness value of 1 indicates that all taxa within a genus recruited an equal proportion of reads, while an evenness of close to zero indicates that only a single taxon recruited most of the reads.

**Figure S4:**
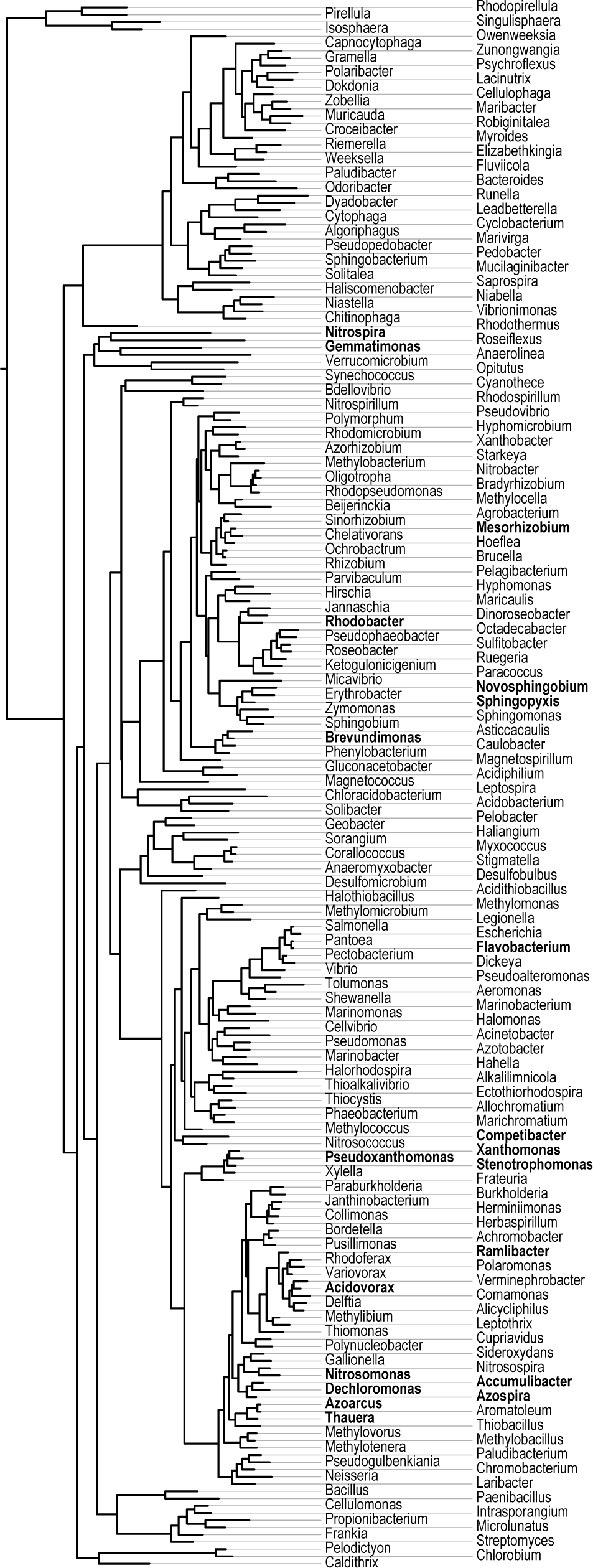
Phylogenetic tree of the genera occurring on granules. The tree is based on 16S sequences from a single representative species of each genus, respectively. The top 20 variance explaining genera are indicated in bold.

**Figure S5:**
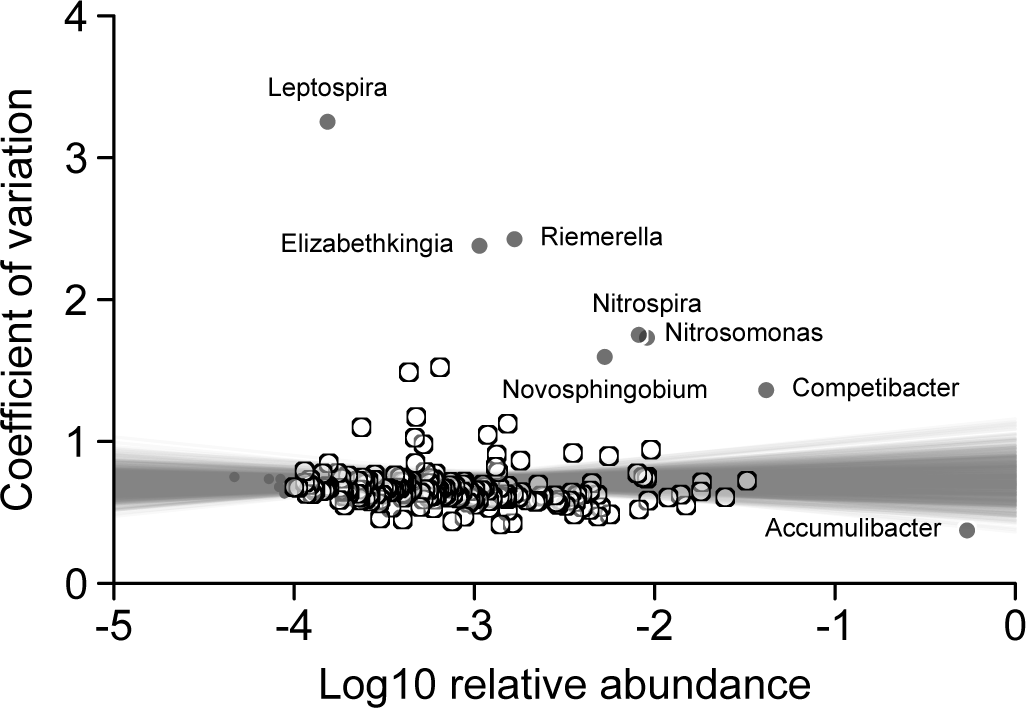
A subset of genera vary more strongly than the mean genus. Each points shows the mean relative abundance and coefficient of variation of a genus across all granules. The straight grey lines show the fit of linear regressions across 1000 bootstraps. For each bootstrap replicate, genera were resampled with replacement. Extremely rare genera with a mean relative abundance of below 10^−4^ were not included in the regression model. Genera that had a Cook’s distance above or below the mean of two times the standard deviation are colored in grey and labeled.

**Figure S6:**
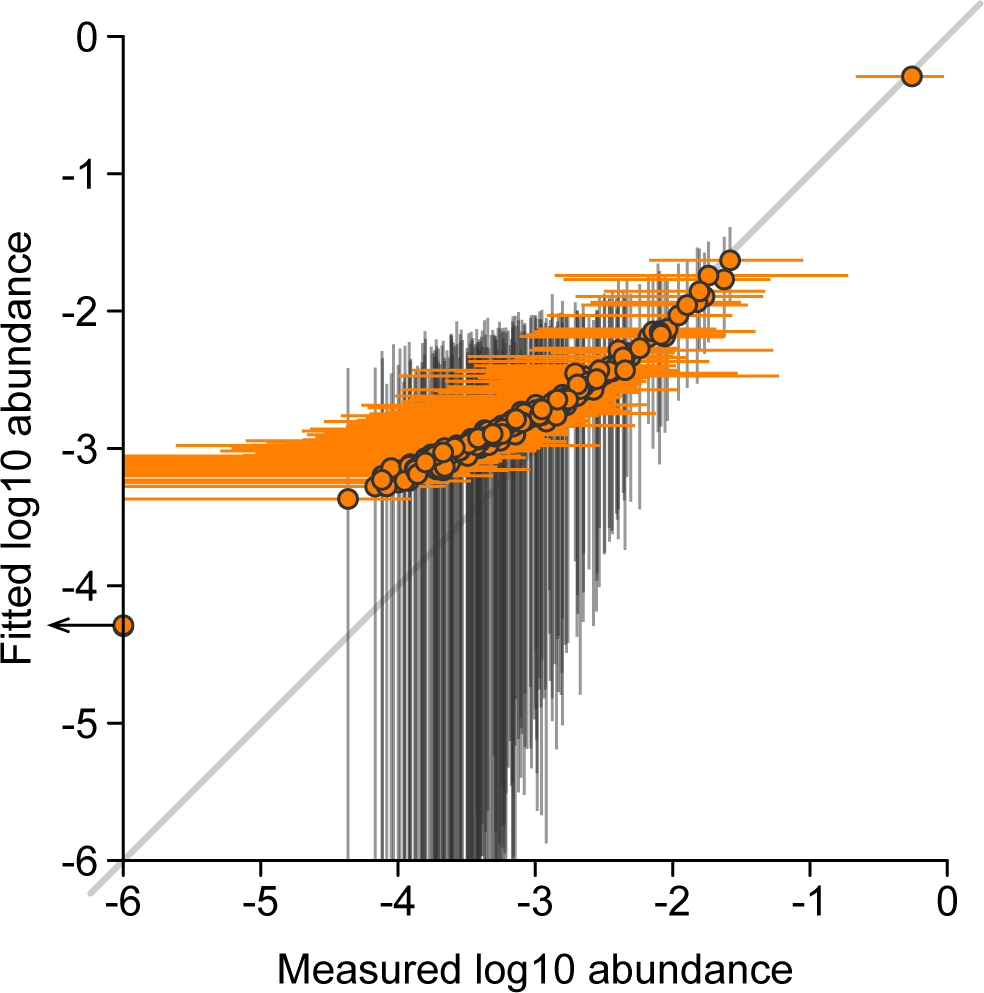
Observed versus fitted relative abundances under the Sloan neutral model. The circles show the maximum likelihood relative abundances of each genus in the metacommunity as a function of the median relative abundance of the genus across the granules. Orange horizontal bars show the actual observed 95% quantile of relative abundance across the granules, and black vertical bars show the 95% quantile for data simulated under the maximum likelihood neutral model with *Nm* = 258.4.

**Figure S7:**
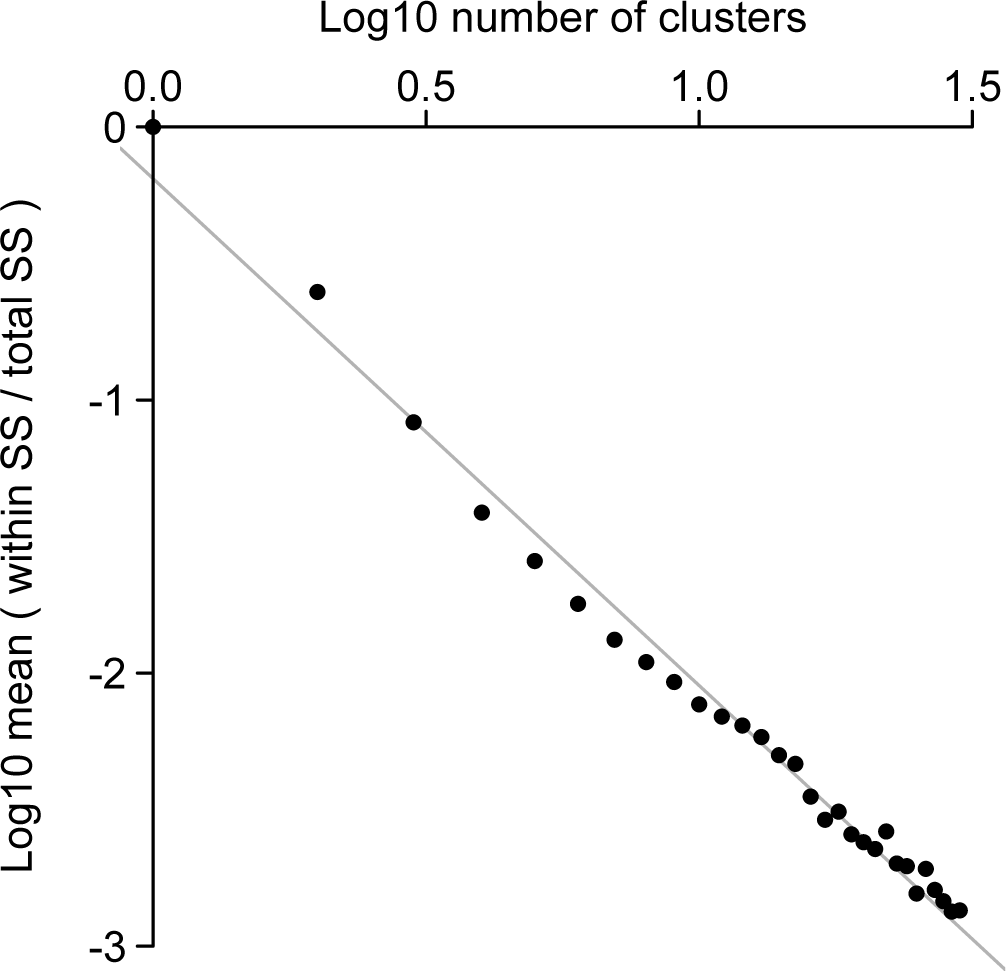
k-means clustering of the log relative abundances. The within versus between cluster sum of squares decreases as a power-law with the number of clusters (exponent ≈ –1.56). Hence, there does not appear to be a concrete cutoff for the best number of clusters.

**Figure S8:**
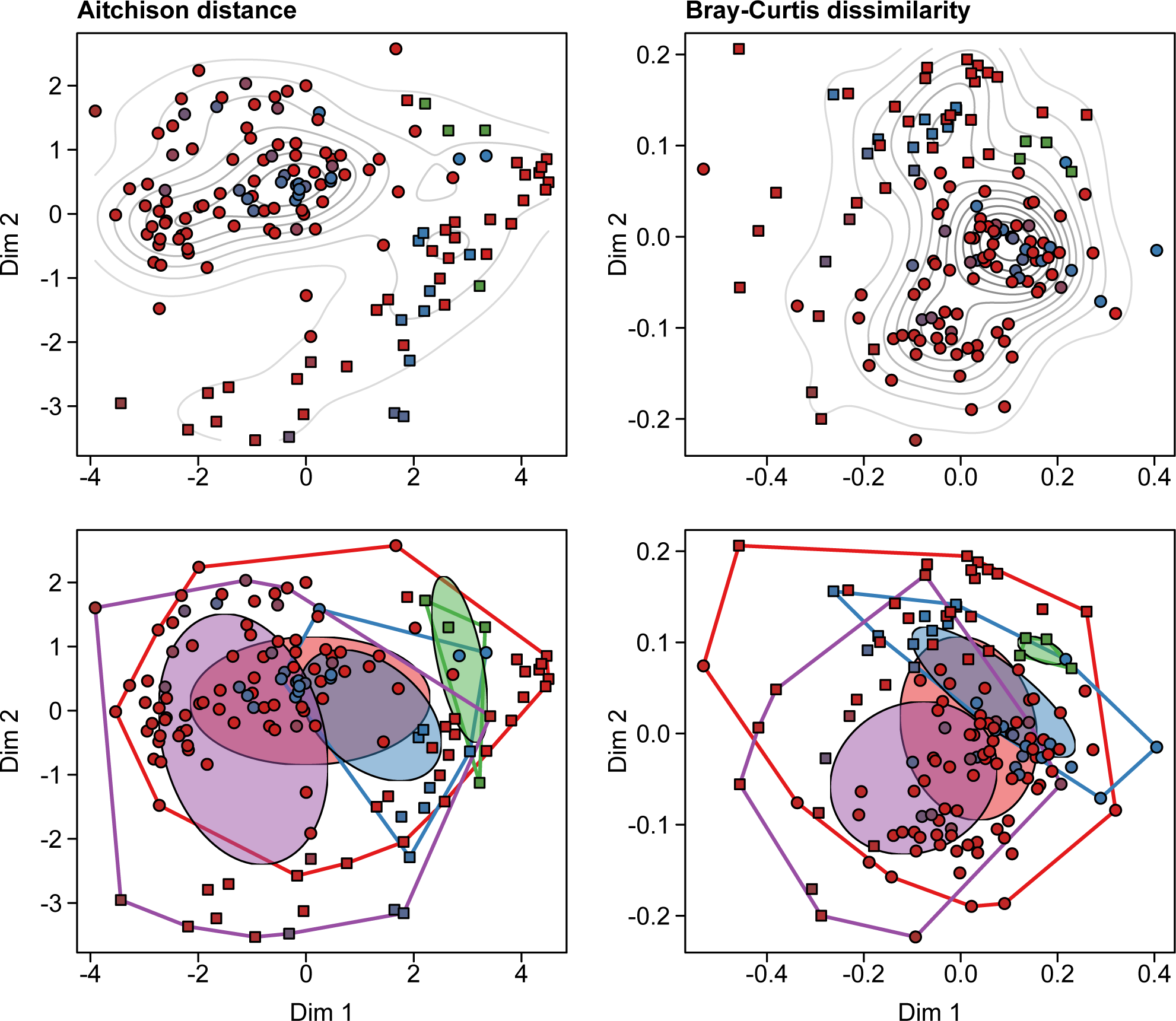
Two dimensional ordination plots based on multi-dimensional scaling. The mapping dimension for the Aitchison and BrayCurtis distance is set to three and two, respectively. The bottom panels show the same layout as the top, but highlights the estimated ellipses and hulls.

**Figure S9:**
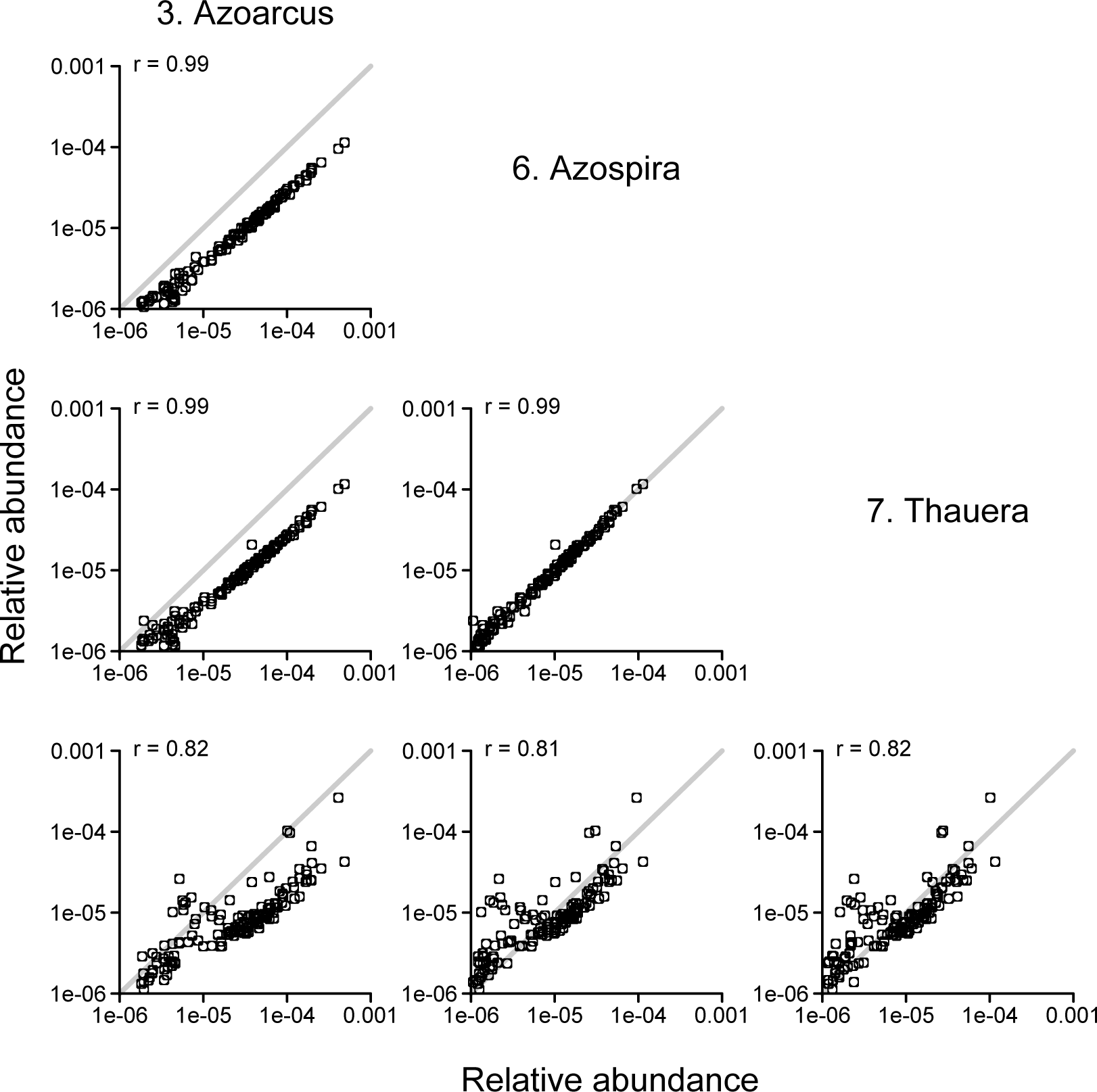
Covariation for a subgroup of genera.

**Figure S10:**
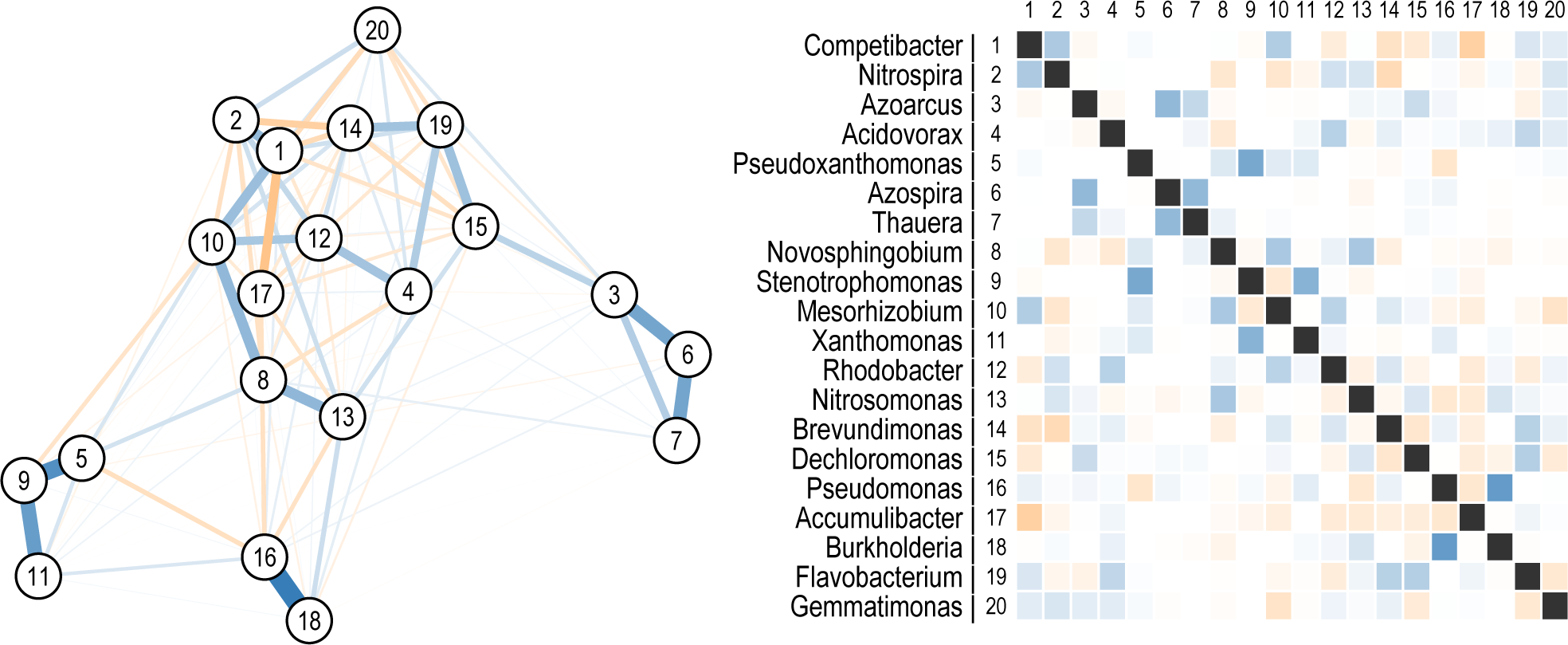
Graphical representation of the inverse covariance matrix for the top 20 varying genera. Blue and orange edges/boxes indicate positive and negative conditional dependencies between genera, respectively. Edge thickness and color intensity indicate the strength of the conditional dependence. The inverse covariance matrix for all genera was calculated using the graphical Lasso approach^58^, with a regularization parameter of *ρ* = 10^−4^.

**Figure S11:**
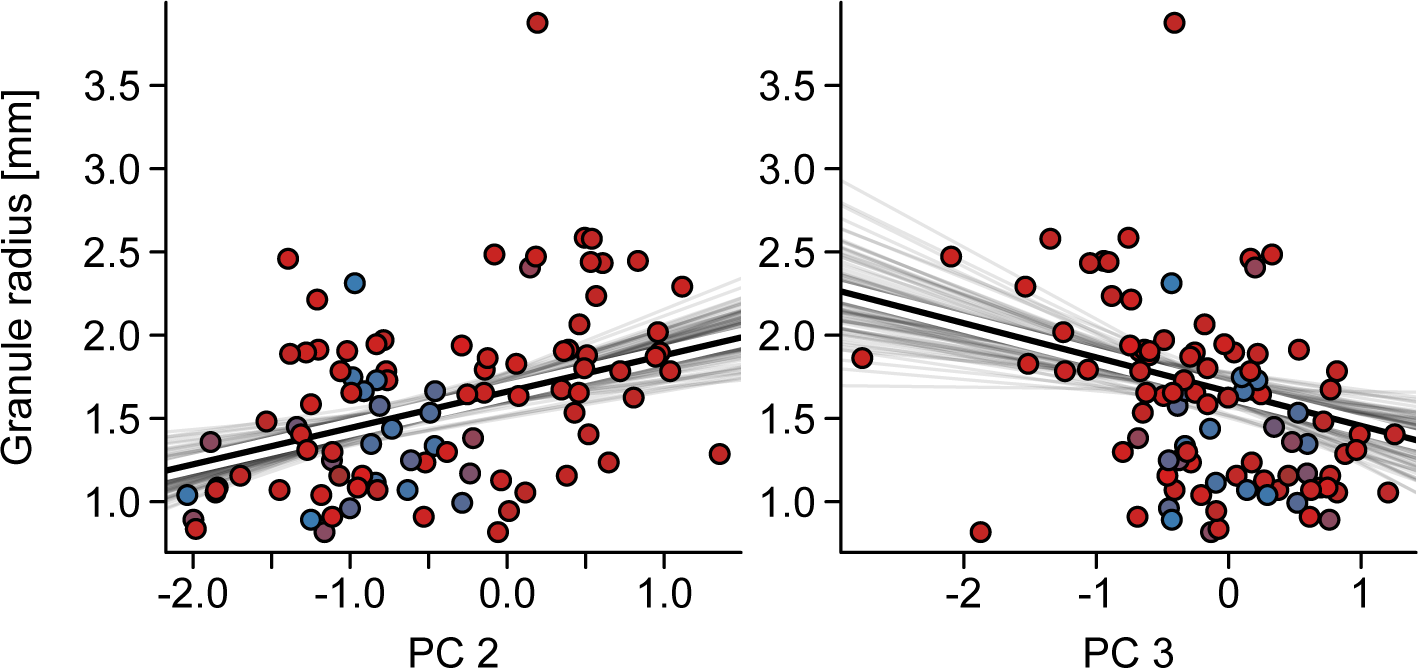
Linear regression of granule radius on principal components two and three.

**Figure S12:**
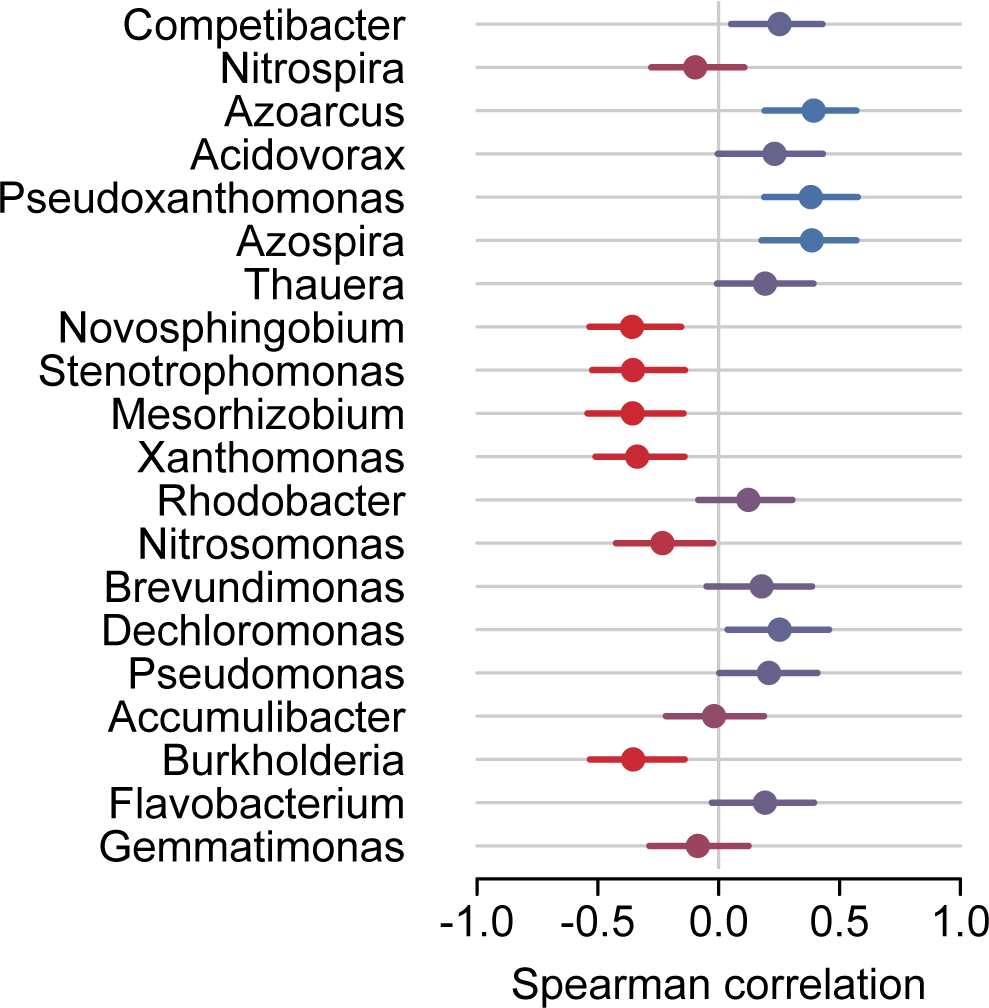
Genera differ in relative abundance between small and large granules. The points show the Spearman correlation between granule radius and relative abundance. Bootstrap error bars were determined through 1000-fold resampling of granules and recalculation the Spearman correlation coefficient.

**Figure S13:**
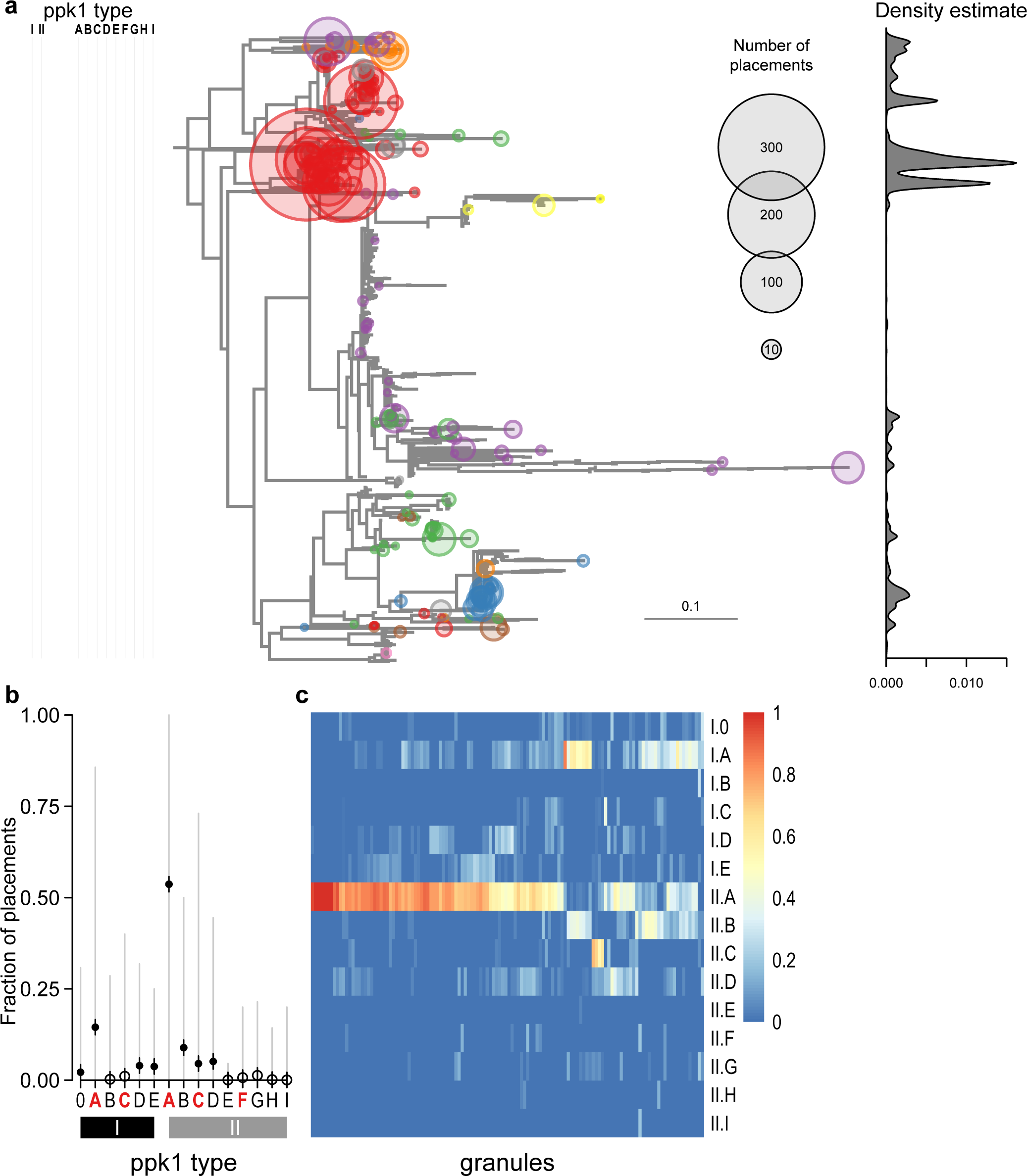
Accumulibacter *ppk1* analysis.

**Figure S14:**
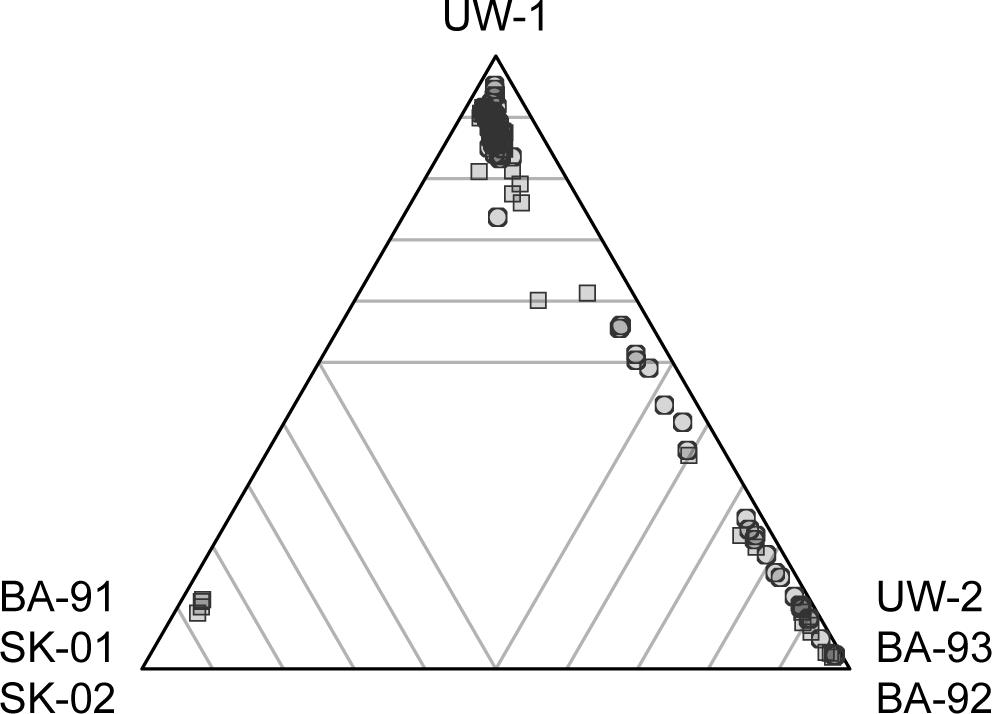
Most granules are dominated by a single Accumulibacter clade. The position on the simplex shows the relative abundance of the three dominant Accumulibacter clades in each granule. The corners correspond to 100% of a single clade. The grey lines show relative abundances of 90%, 80%, 70%, 60%, and 50% moving from a corner to the center of the simplex.

**Figure S15:**
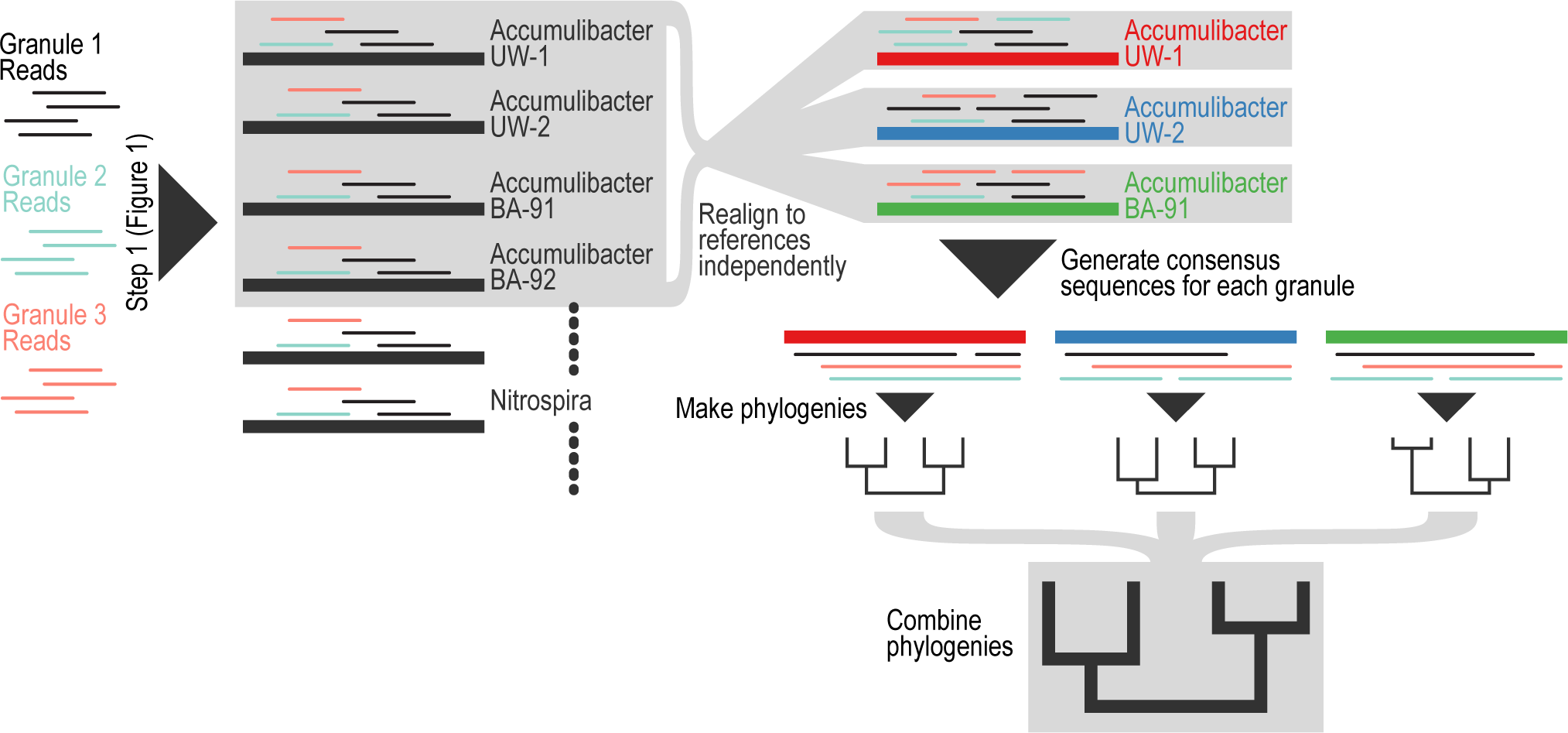
Methodological overview of phylogenetic reconstruction from reference genomes using the short reads.

**Figure S16:**
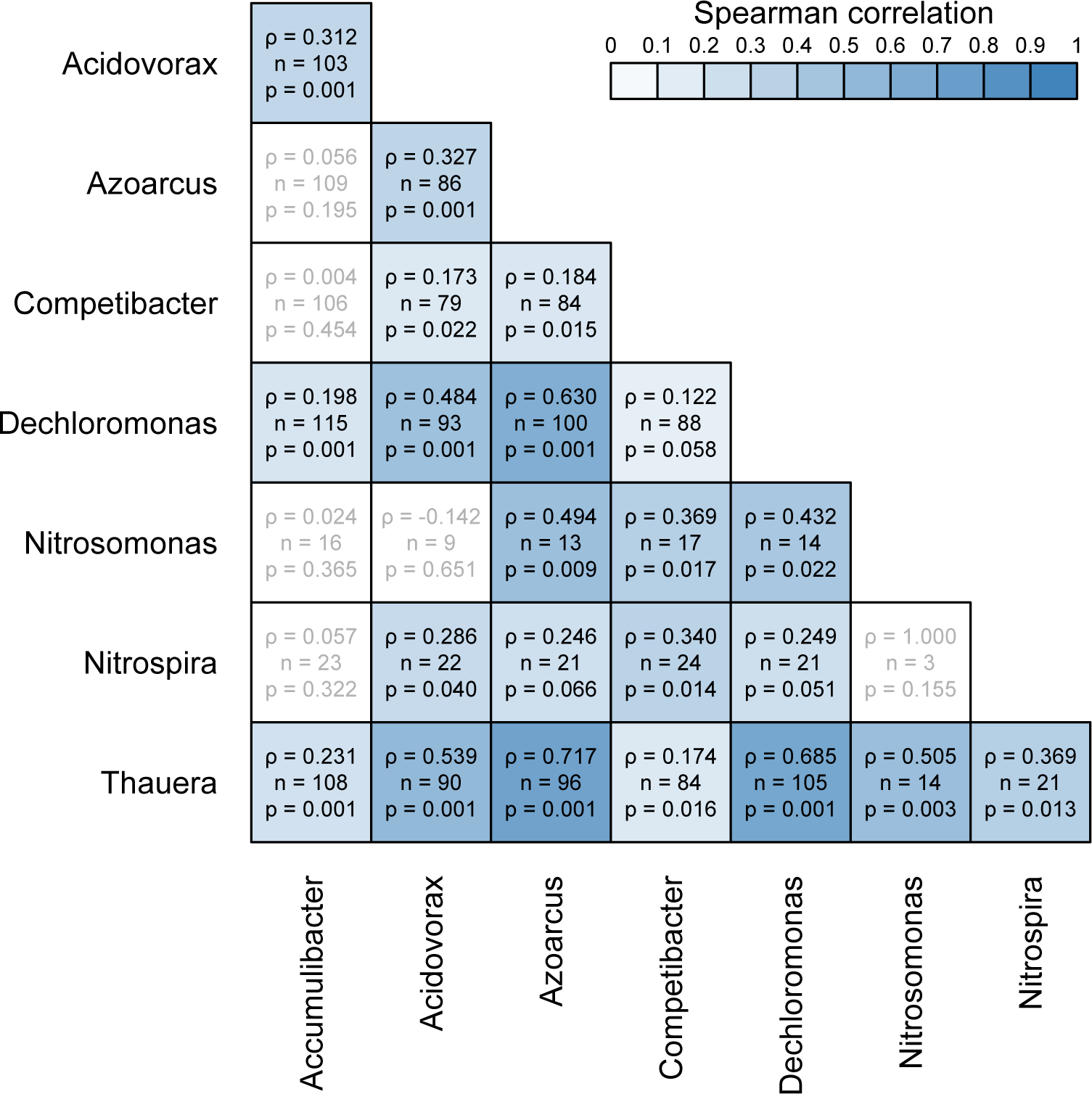
Spearman correlation between cophenetic distances of granules based on different genera. Statistically significant correlations (*ρ* > 0 with *p* < 0.1) are in black type, while non-significant correlations are in grey type. Each square shows Spearman’s rank correlation *ρ*, the number of granules for which there was enough coverage in both genera to be included in the analysis, and the empirical *p*-value from a Mantel test.

**Figure S17:**
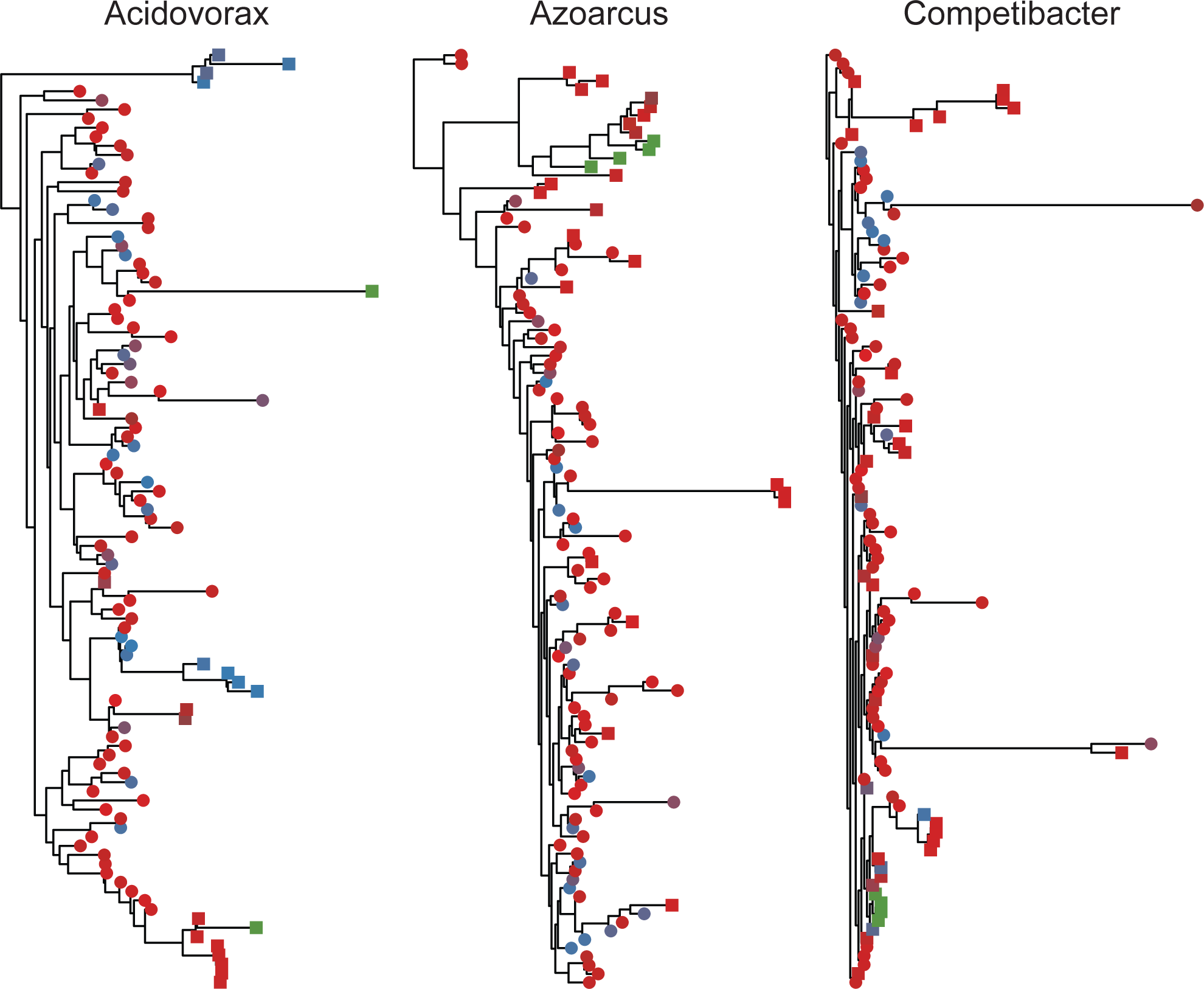
Granule phylogenies for Acidovorax, Azoarcus, and Competibacter.

**Figure S18:**
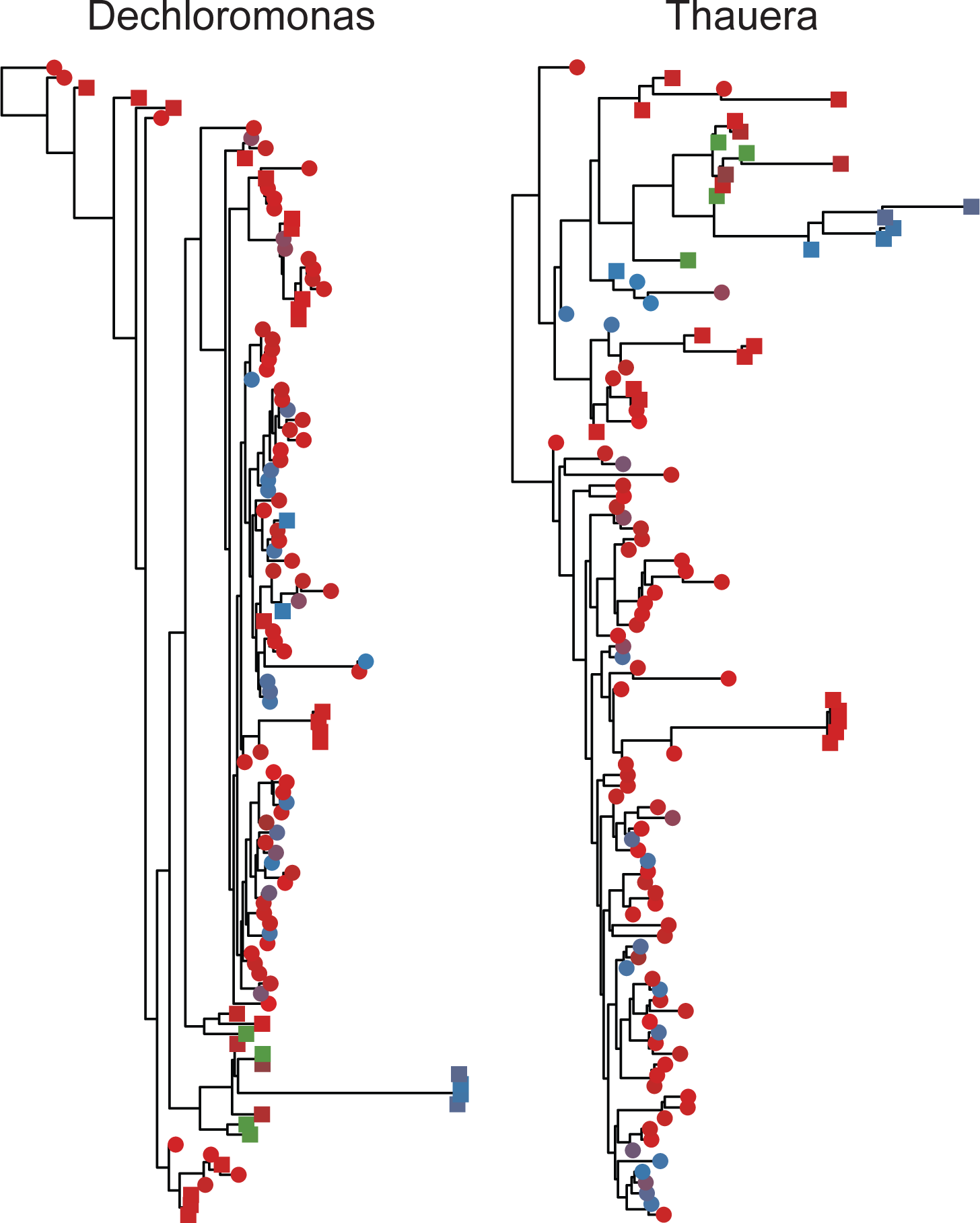
Granule phylogenies for Dechloromonas and Thauera.

**Figure S19:**
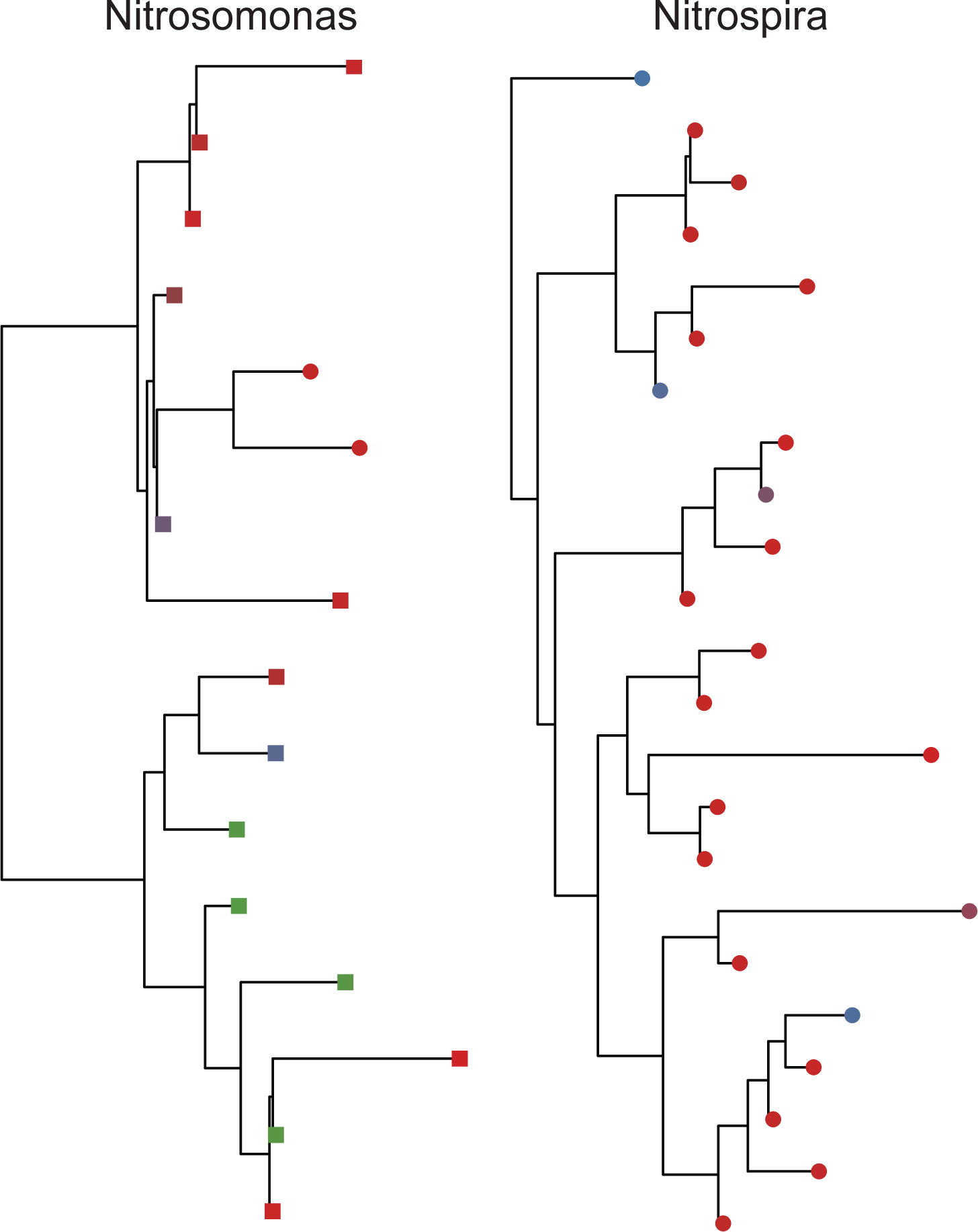
Granule phylogenies for Nitrosomonas and Nitrospira.

